# Folate depletion induces erythroid differentiation through perturbation of de novo purine synthesis

**DOI:** 10.1101/2022.12.22.521608

**Authors:** Adam G. Maynard, Nancy K. Pohl, Boryana Petrova, Alan Y.L. Wong, Peng Wang, Andrew J. Culhane, Leah Hirsch, Ngoc Hoang, Orville Kirkland, Tatum Braun, Sarah Ducamp, Mark D. Fleming, Hojun Li, Naama Kanarek

## Abstract

All dividing cells require the essential vitamin folate. Hematopoietic cells harbor a unique sensitivity to folate deprivation, as implied by the development of folate-deficient anemia and the utility of anti-folate chemotherapy in blood cancer. To study this metabolic sensitivity, we applied mild folate depletion to human and mouse erythroid cell lines, as well as primary murine erythroid progenitors. We show that folate depletion induces early blockade of purine synthesis that is followed by enhanced heme metabolism, hemoglobin synthesis and erythroid differentiation. This finding is phenocopied by inhibition of folate metabolism using SHIN1, an inhibitor of the folate enzymes SHMT1/2. The metabolically-driven differentiation is rescued by supplementation of purine precursors, yet occurs independent of nucleotide sensing through mTORC1 and AMPK. Our work profiles the metabolic response to folate depletion in erythroid cells and suggest that premature differentiation of folate-deprived erythroid progenitor cells is a mechanistic etiology to folate-deficiency induced anemia.

## Introduction

Folic Acid (FA) is a vitamin required for many cellular processes, including nucleotide synthesis^1^, methylation^2^, and redox balance^3^, collectively known as one-carbon (1C) metabolism^4^. 1C metabolism requires the folate-mediated transfer of methyl or formyl groups from 1C donors, such as serine, to 1C acceptors, such as purine nucleotide intermediates. Folate’s role as the 1C unit carrier is essential in all cells; however, folate is not synthesized by mammalian cells and must be consumed through the diet. Folate deficiency affects all cell types, but in post-embryonic tissues is especially detrimental to hematopoietic cells. This sensitivity is demonstrated by the primary manifestation of nutritional folate deprivation - anemia^5^.

Erythroid differentiation, or erythropoiesis, is the process that generates red blood cells from pluripotent hematopoietic cells in the bone marrow. During this multistep process, progenitor cells go through several proliferation cycles to achieve abundant cell counts while they also decrease in size, increase hemoglobin synthesis, and undergo nuclear ejection on their way to form mature erythrocytes. A notable metabolic change during erythropoiesis is the upregulation of heme synthesis. Heme synthesis initiates with condensation of mitochondrial glycine with Succinyl-CoA by the rate limiting enzyme, 5-aminolevulinic acid synthase 2 (ALAS2) to generate heme in an 8-step reaction. The key metabolite, glycine, can be generated from serine via 1C metabolism, or through transport of extracellular glycine.

The connection between folate and erythroid differentiation was revealed in the 1940s when megaloblastic anemia was treated with folate supplementation^5^. Early studies that sought to understand the progression of folate-deficient anemia suggested DNA damage-induced apoptosis as the mechanism for reduced erythrocytic cell counts^6,7^. However, this suggested mechanism does not fully support the appearance of large, hemoglobin-rich, red blood cells that are characteristic of megaloblastic anemia. This leaves the molecular etiology of nutritional megaloblastic anemia un-resolved.

We investigated the molecular etiology of nutritional anemia by studying the cellular consequences of mild folate depletion in erythroid cells. We show that purine synthesis is inhibited early after a drop in folate availability, and that this is followed by the induction of an erythroid differentiation program. Bypassing de novo purine synthesis by supplementation of purine precursors relieved this aberrant differentiation. Folate-deprivation induced erythroid differentiation is phenocopied by pharmacological inhibition of folate metabolism using the SHMT1/2 enzymes inhibitor SHIN1, and occurs also in primary erythroid progenitor cells derived from murine fetal livers. Early, aberrant progenitor differentiation can result in low red blood cells counts due to reduced number of proliferation cycles, and can also explain the appearance of large, heme-filled erythrocytes, both are characteristics of megaloblastic anemia, the type of anemia caused by folate deprivation.

## Results

### Folate depletion upregulates heme synthesis and induces erythroid differentiation

To investigate the metabolic changes in erythroid cells following a mild folate depletion that might be similar to the stress induced by nutritional folate deprivation, we cultured erythroid lineage cells in 100 nM FA, a concentration at least 20-fold lower than standard culture media. This concentration allows cell proliferation in culture, but induces a mild proliferation defect (Fig. 1a). 100 nM FA is a similar magnitude to the plasma folate concentration in healthy adults (~20-80 nM)^8,9^ as opposed to the supraphysiological FA concentration in standard culture media (RPMI – 2,000 nM or DMEM – 9,000 nM)^10^. Through metabolite profiling of 6-day folate-deprived K562, a chronic myelogenous leukemia cell line with the potential to undergo erythroid differentiation^11,12^, we observed significant accumulation of the purine synthesis intermediates 5’-phosphoribosyl-glycinamide (GAR) and 5’-phosphoribosyl-5-aminoimidazole-4-carboxamide (AICAR) (Fig. 1b). These two metabolites are substrates for reactions that require 10-formyl tetrahydrofolate (10-formyl THF) and are expected to become the rate limiting steps of de novo purine synthesis upon folate depletion^13^. Several nucleotide salvage intermediates, including hypoxanthine, inosine, and guanosine, are depleted under low FA conditions (Fig. 1b), as these can be used to regenerate inosine monophosphate (IMP) levels when purine synthesis is impaired^14^. Unexpectedly, we also observe a significant increase in intracellular heme, as well as accumulation of the last intermediate of heme synthesis, protoporphyrin IX (Fig. 1b, c). To test whether heme synthesis is actively enhanced in folate-depleted K562 cells, we assayed the mRNA expression of heme biosynthesis pathway genes. We found significant upregulation of several heme biosynthesis genes (Fig. 1d), including the rate-limiting enzyme ALAS2^15^. While depletion of folate increases mRNA expression of heme biosynthesis pathway components, there is no accompanied upregulation of 1C metabolism genes (Extended Data Fig. 1a). The synthesis of a single heme molecule requires 8 molecules of glycine that can be generated from serine through the action of the folate-dependent enzyme SHMT, or imported from exogenous pools. Although serine was underutilized in low FA conditions (Extended Data Fig. 1b), as expected from folate depletion, glycine levels were still sufficient to support heme synthesis due to utilization of exogenous glycine (Extended Data Fig. 1c). The increase in total heme was not associated with an increase in mitochondrial mass or membrane potential, which suggests a utilization of heme outside of the mitochondrial respiratory complexes (Extended Data Fig. 1d, e).

**Figure 1:**
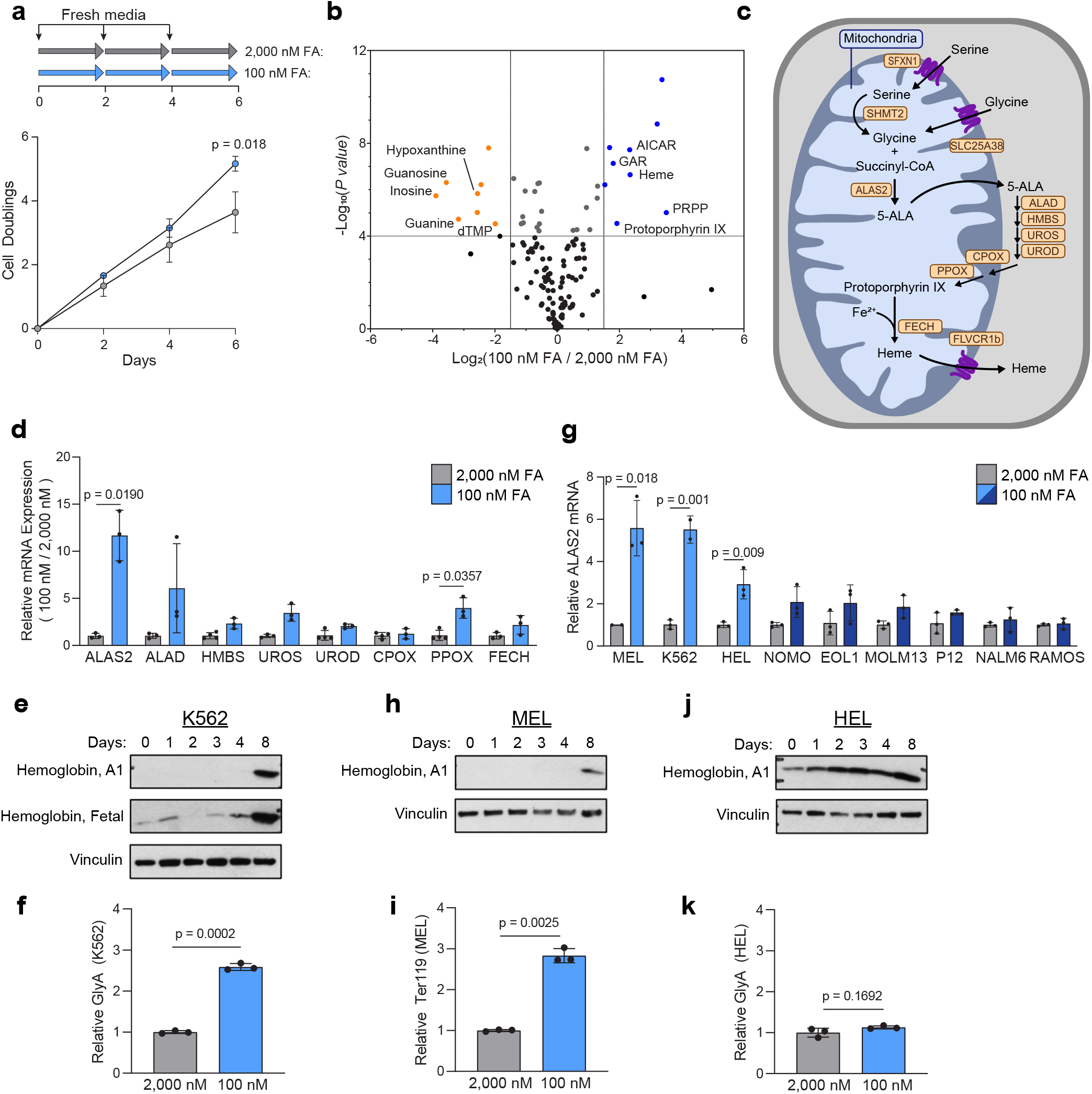
Folate depletion induces heme synthesis and differentiation in erythroid cell lines. **a,** Experimental timeline (top) and proliferation rate (bottom) of K562 cultured for 6 days in folate-free RPMI supplemented with 2,000 or 100 nM Folic Acid (FA). Media was changed every 2 days. **b,** Volcano plot depicting day 6 metabolic changes between K562 cells cultured in 2,000 and 100 nM FA. Metabolites’ abundance was measured by liquid chromatography-mass spectrometry (LC-MS). **c,** Schematic of the mammalian heme biosynthesis pathway. **d,** RT-qPCR analysis of heme biosynthesis genes in 2,000 nM FA (grey) vs. 100 nM FA (blue). **e,** Western Blot analysis of hemoglobin A1 and fetal expression in K562 over 8 days in 100 nM FA. Vinculin is a loading control. **f,** Cell surface Glycophorin A (GlyA) levels in K562 measured by flow cytometry. **g,** RT-qPCR analysis of ALAS2 in the indicated cell lines following 6 days culture in 2,000 or 100 nM FA. (light blue = erythroid, dark blue = non-erythroid) **h,** Western blot analysis of hemoglobin A1 in MEL. Vinculin is a loading control. **i,** Cell surface Ter119 levels in MEL measured by flow cytometry. **j,** Western Blot analysis of hemoglobin A1 in HEL. Vinculin is a loading control. **k,** Cell surface Glycophorin A (GlyA) levels in HEL measured by flow cytometry. Data shown are mean (± s.d.) of three biological replicates. P values were calculated using an unpaired Student’s t-test.

Increased heme synthesis is a hallmark of the erythroid differentiation process^16^. As K562 cells have the potential to undergo erythroid differentiation, we investigated whether low FA induces erythroid differentiation in these cells. Over 8 days in low FA, fetal hemoglobin (found in K562 cells even though these are not embryonic-derived cells^17^), and hemoglobin A1 (Hba1) levels increase (Fig. 1e), as well as the erythroid cell surface marker Glycophorin A (Fig. 1f). These data suggest erythroid differentiation of folate-deprived K562s.

To test if folate deprivation induces erythroid differentiation in additional hematopoietic cell lines, we measured ALAS2 induction in 9 hematopoietic cell lines following folate deprivation. We observed upregulation of ALAS2 only in the 3 erythroid leukemia lines included in the panel (MEL, K562, and HEL), and not in cell lines derived from other lineages (Fig. 1g). The induction of ALAS2 expression was not correlated with growth rate (Extended Data Fig. 1f). Additionally, we observed no induction of the non-erythroid heme-synthesis enzyme, ALAS1 (Extended Data Fig. 1g). This supports an erythroid specific role of heme synthesis in folate-deprived cells as opposed to a broad upregulation of heme synthesis in all leukemia cell lines. We therefore validated the induction of heme synthesis and the increase in differentiation markers following mild folate deprivation in two additional erythroid cell lines; the mouse line, MEL, and the human line, HEL. Both MEL and HEL increased hemoglobin A1 protein expression (Fig. 1h, j). We assayed the erythroid differentiation surface markers Ter119 and Glycophorin A found on MEL and HEL, respectively, following folate deprivation. While no change in Glycophorin A surface expression was observed in HEL, MEL significantly increased expression of Ter119 (Fig. 1i, k). Taken together, our data reveals induction of erythroid differentiation by mild folate deprivation.

Our metabolite profiling data of 6-day folate-deprived K562 revealed accumulation of protoporphyrin IX, a metabolite that, together with iron, generates a complete heme molecule (Fig. 1c and Extended Data Fig. 2a). This raised the possibility that folate-deprived K562 suffer from lack of iron due to increased demand from upregulated heme synthesis. However, neither iron supplementation, either as free iron (Fe^2+^) or transferrin-bound iron (Fe^2+^-transferrin), nor iron chelation (using DFO), altered proliferation (Extended Data Fig. 2b), or differentiation (Extended Data Fig. 2c, d), although these treatments result in the expected changes in transferrin receptor gene expression (Extended Data Fig. 2e). These data rule out a role for iron availability or limitation as a driver of folate-deprivation induced differentiation.

### Purine biosynthesis inhibition is an early event following folate deprivation and purine supplementation rescues folate depletion-induced differentiation

FA is directly and indirectly involved in many metabolic reactions, including nucleotide synthesis, amino acid metabolism, and methylation reactions. Therefore, we asked which metabolic changes in folate-deprived erythroid cells drive differentiation. We addressed this by first mapping the kinetics of the cellular response to low folate availability (Extended Data Fig. 3a); Transcriptionally, induction of heme biosynthesis gene expression in K562 occurs at day 4 following folate deprivation (Fig. 2a). Heme levels were increased later than these gene expression changes, at day 6 (Fig. 2b). Metabolically, the purine synthesis pathway is the first pathway to show significant perturbations in both K562 and MEL cells as early as day 2 following folate deprivation: at day 2, K562 and MEL cells had 7 and 21 significantly altered metabolites, respectively, with AICAR, GAR and Ribose-1-phosphate (R1P) the only shared metabolites between the two cell lines (Fig. 2c, d). AICAR and GAR are intermediates of the purine synthesis pathway and R1P is the byproduct of purine salvage reactions (Extended Data Fig. 3b). Although folate is essential for IMP and dTMP synthesis, examination of nucleotide levels in K562 and MEL showed that while there is a slight reduction in ATP and GTP in K562 and MEL over 4 days of FA depletion, no major pyrimidine synthesis inhibition occurs (Extended Data Fig. 3c, d). Therefore, although pyrimidine synthesis inhibition can be associated with myeloid leukemia differentiation^18^, it is not likely to be the metabolic driver of the erythroid differentiation we observe here.

**Figure 2:**
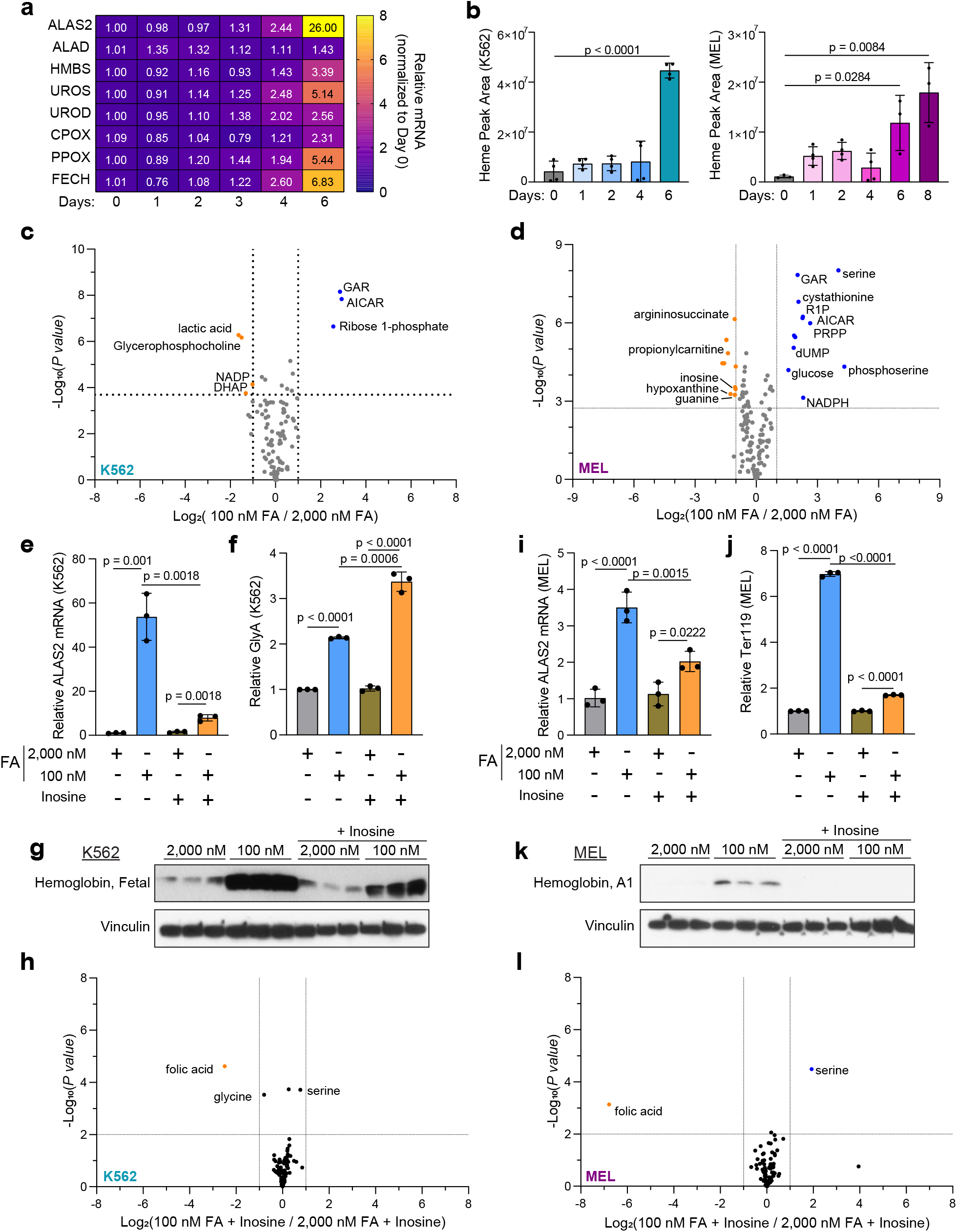
Purine synthesis is disrupted early following folate deprivation, and this disruption is essential for folate-deprivation induced differentiation. **a,** RT-qPCR analysis of heme synthesis genes in K562 over 6 days in 100 nM FA. **b,** Total intracellular heme in K562 (left) and MEL (right), measured by LC-MS. **c, d,** Volcano plots depicting the metabolic changes between 2,000 and 100 nM FA at day 2 in K562 (**c**) and MEL (**d**). **e,** RT-qPCR analysis of ALAS2 mRNA levels in K562 following 6 days in 2,000 or 100 nM FA, with or without inosine (100 μM) supplementation. **f,** Cell surface GlycophorinA (GlyA) in K562 following 6 days in 2,000 or 100 nM FA with or without inosine (100 μM) supplementation measured by flow cytometry. **g,** Western blot analysis of hemoglobin levels following 6 days in 2,000 or 100 nM FA with or without inosine supplementation (100 μM). Vinculin is a loading control. **h,** Metabolic changes in K562 cultured in inosine supplemented 100 nM FA vs 2,000 nM FA media. **i – l,** The same experiments as 2e-f in MEL. Data shown are mean (± s.d.) of three biological replicates. All P values were calculated using an unpaired Student’s t-test.

Previous studies in erythroleukemia have shown that both purine supplementation^19,20^ and inhibition of purine synthesis^21,22^ can induce erythroid differentiation. Therefore, the early inhibition of purine synthesis led us to hypothesize that perturbation of this pathway drives a stress response that results in erythroid cell differentiation. To assess this hypothesis, we rescued purine synthesis by supplementation with inosine (100 μM), a purine salvage intermediate, that enables purine synthesis while bypassing folate-dependent de novo synthesis. Inosine supplementation restored the proliferation rate of K562 and MEL cells in low FA (Extended Data Fig. 3e, f). There was no effect of inosine on proliferation in high FA cultured cells. Inosine supplementation also blocked folate-deprivation induced differentiation, as observed by the lack of induction of ALAS2 expression (Fig. 2e, i), the reduction in hemoglobin synthesis (Fig. 2g, k), and the reduction in the erythroid surface marker Ter119 in MEL cells (Fig. 2j, Extended Data Figure 4a-c). However, GlyA surface expression was not rescued with inosine in K562 (Fig. 2f, Extended Data Figure 4d-f). In addition, treatment with inosine erased nearly all the metabolic changes associated with low FA at day 2 in K562 and MEL cells; only FA, serine, and glycine were significantly different (Fig. 2h, l). As expected, inosine treatment rescued de novo purine synthesis defects and nucleotide levels (Extended Data Fig. 3g-j). This suggests that neither the absence of FA, nor the accumulation of intracellular serine, drive differentiation on their own. These data may also indicate that inosine supplementation in low FA conditions allows limiting pools of folate molecules to be redirected to other non-purine, folate-dependent reactions. Low FA media supplemented with inosine may permit cells to maintain all cellular folate-required reactions and avoid the metabolic stress that is otherwise associated with folate deprivation. Further work will be needed to address this specific question.

### mTORC1 inactivation and AMPK activation are not sufficient to induce erythroid differentiation

The early metabolic changes following folate deprivation include an increase in AICAR levels and decrease in purine levels. High levels of AICAR activate AMPK signaling^23^, and depletion of purines inhibits mTORC1 activation^24^ (Fig. 3a). We therefore tested the hypothesis that either one, or both, of these major signaling pathways drive the differentiation program.

**Figure 3:**
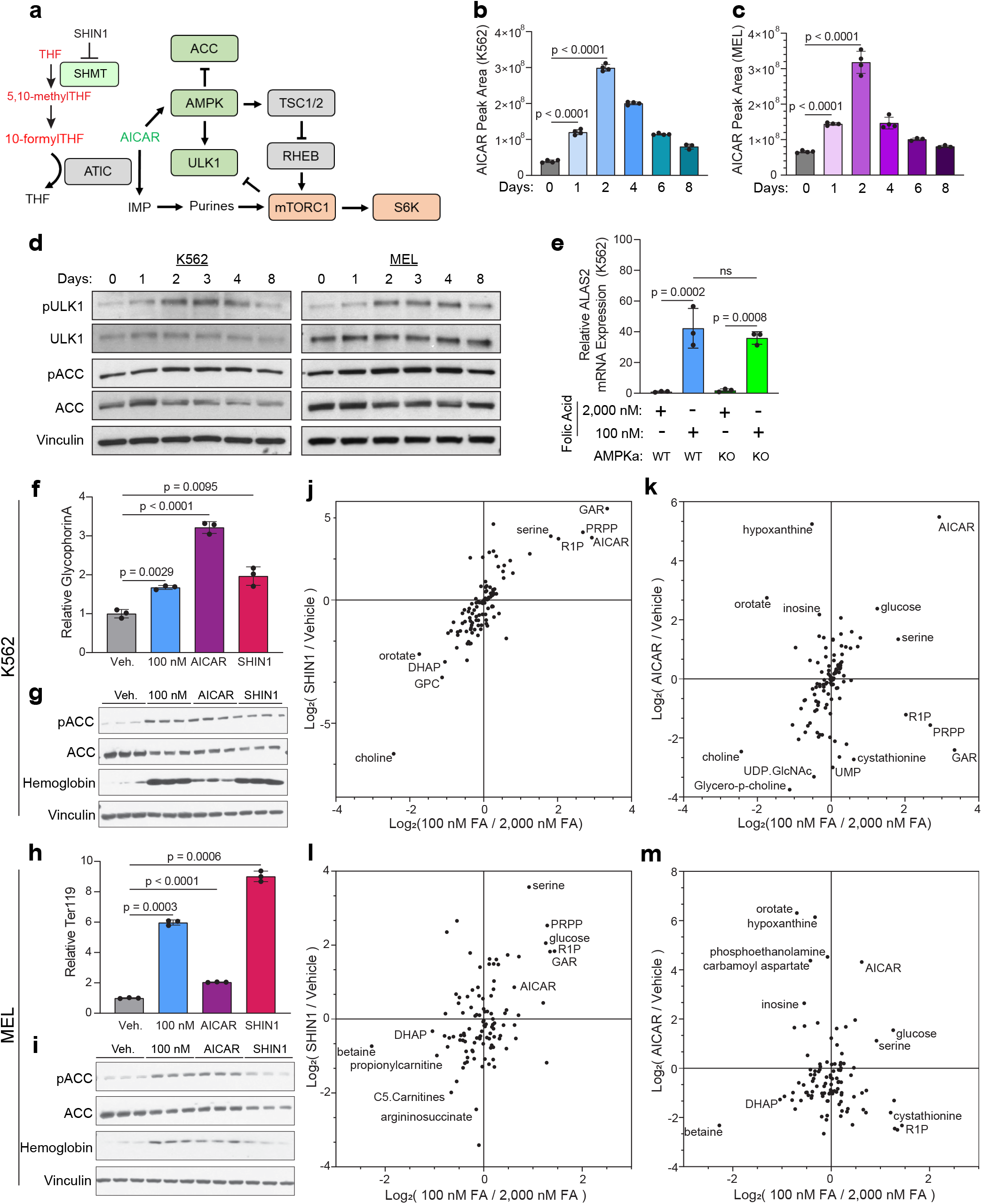
AICAR, but not AMPK activation, is sufficient to induce folate-deprivation induced differentiation. **a,** Schematic depicting the connection between folate depletion, purine synthesis, and AMPK and mTORC1 signaling. **b, c,** AICAR levels in K562 (**b**) and MEL (**c**) over 8 days in 100 nM FA measured by LC-MS. **d,** Western blot analysis of AMPK signaling in K562 (left) and MEL (right) over 8 days culture in 100 nM FA. Vinculin is a loading control. **e,** RT-qPCR analysis of ALAS2 mRNA expression in AMPKa1/a2 WT and double knockout (KO) K562 cells cultured in 2,000 or 100 nM FA for 6 days. **f,** Cell surface Glycophorin A expression measured by flow cytometry in K562 cultured in 100 nM FA, or treated with SHIN1 (1.25 μM), or AICAR (500 μM) for 2 days. Vehicle, AICAR, and SHIN1 were all cultured in 2,000 nM FA. (Veh – vehicle). **g,** Western blot analysis of phosphorylated ACC (pACC)/ACC and hemoglobin levels in K562 in the indicated treatments, as in f Vinculin is a loading control. **h, i,** Analysis of cell surface Ter119 (**h**) and AMPK signaling (**i**) at Day 2 in the indicated treatments, as in f, g. **j-m,** comparison of the fold change (Log2) of 100 nM/2,000 nM FA vs SHIN1/Vehicle (**j, l**) and AICAR/Vehicle vs 100 nM/2,000 nM (**k, m**), in K562 (**j, k**) and MEL (**l, m**). Data shown are mean (± s.d.) of three biological replicates. All P values were calculated using an unpaired Student’s t-test.

While purines were significantly depleted in K562 cells at early timepoints (Extended Data Fig. 3c) a closer look at the magnitude of purine depletion revealed only a mild depletion in MEL at the early time points (Extended Data Fig. 3d). Accordingly, we observed a decrease in mTORC1 activation in K562, but to a lesser extent in MEL (Extended Data Figure 5a). Further, perturbation of the mTORC1 signaling pathway by the inhibitors Torin1 and Rapamycin in high FA conditions did not induce differentiation of K562 and MEL cells (Extended Data Fig. 5b-e), ruling out sufficiency of mTORC1 inactivation for driving folate-deprivation induced erythroid differentiation.

Unlike the somewhat inconsistent purine depletion between the two cell lines, the levels of the AMPK activator AICAR (Fig. 3a) increase significantly as early as one day following folate deprivation in both K562 and MEL cells (Fig. 3b, c). AICAR accumulation, as well as other intermediate metabolites of the purine synthesis pathway (Extended Data Fig. 6a, b), occurs early enough to be acting as an inducer of the differentiation program. Indeed, AMPK signaling is activated shortly after the elevation in AICAR levels as measured by the phosphorylation of ACC and ULK1, two principle downstream targets of AMPK^25^ (Fig. 3d). To investigate whether AMPK signaling is sufficient to induce erythroid differentiation we applied the pharmacologic AMPK activator GSK621^26^. While this compound does not compromise the proliferation rate of K562 and MEL cells to the extent observed in cells cultured in 100 nM FA (Extended Data Fig. 6c, d), and it activates AMPK (Extended Data Fig. 6e, f), it does not induce differentiation in K562 and MEL cells (Extended Data Fig. 6g, h). Further, when we tested whether AMPK signaling is necessary for folate-depletion induced differentiation by genetically targeting both AMPKa1 and AMPKa2 (AMPKa double knockout - DKO) (Extended Data Fig. 6i, j), we observed no blunting of ALAS2 expression following folate deprivation (Fig. 3e), indicating AMPK signaling is not required for induction of the differentiation program.

### Excess AICAR is sufficient to induce erythroid differentiation

We next wondered if it is possible that AICAR functions independently of AMPK and is sufficient for the induction of erythroid differentiation. We therefore applied two additional approaches to increase AICAR levels: AICAR supplementation, and the 1C metabolism inhibitor, SHIN1^27^. SHIN1 inhibits the enzymes SHMT1 and SHMT2 and lowers intracellular levels of reduced folate, thereby mimicking nutritional folate deprivation and increasing AICAR levels (Fig. 3a). We used 500 μM AICAR and 1.25 μM SHIN1 - doses that slow proliferation rate, but do not induce a cytostatic or cytotoxic effect (Extended Data Fig. 6k, l). Compared to folate deprivation, AICAR supplementation results in significantly higher levels of intracellular AICAR, while SHIN1 treatment results in comparable AICAR accumulation (Extended Data Fig. 6m, n). Similar to cells cultured in low folate, cells treated with either AICAR or SHIN1 differentiate, as evidenced by upregulation of erythroid cell surface markers and accumulation of hemoglobin (Fig. 3f-i). In K562, each of the perturbations – low FA, AICAR, and SHIN1 – resulted in AMPK activation (Fig. 3g). This is in agreement with the high AICAR levels in all three conditions (Fig. 3j, k), which directly activates AMPKa^23^ (Fig. 3a). Interestingly, in MEL cells low FA and AICAR, but not SHIN1, activate AMPK (Fig. 3i). This can be the result of the accumulation of metabolites other than AICAR in the SHIN1-treated MEL cells; one of these metabolites is glucose - an upstream suppressor of AMPK signaling (Fig. 3l, m). These data are consistent with our previous observations that indicate AMPK activation alone is not sufficient to induce differentiation. Additional cellular events that ensue in conditions of 1C metabolism blockade are therefore necessary for the induction of differentiation. The fact that AICAR supplementation alone induces differentiation is fascinating and implies that the AICAR-induced cellular response likely includes some downstream events that are beyond AMPK activation and that are essential for the metabolically-induced erythroid differentiation.

Our metabolite profiling of K562 and MEL cells treated with low FA, AICAR and SHIN1 revealed a significant reduction in the choline metabolism pathway, including choline, glycerophosphocholine, and phosphocholine (Fig. 3j-m, Extended Data Fig. 7a). The metabolic association between choline and folate metabolism stems from a shared reaction: In the methionine cycle, the folate form 5-methyl THF is required for the conversion of homocysteine to methionine by the enzyme MTHFR. Alternatively, methionine can be synthesized from homocysteine by the enzyme BHMT using betaine, instead of 5-methyl THF, as the methyl donor. Betaine is synthesized from choline by the enzyme CHDH (Extended Data Fig. 7a). This connection presented two possibilities: 1. Choline metabolism is upregulated in low FA to support the methionine cycle and maintain cellular methylation. 2. The depletion of both FA and choline could lead to methylation defects and epigenetic reprogramming that induces differentiation. With these possibilities in mind, and coupled with the fact that choline metabolism is essential for murine and human erythropoiesis^28^, we detected metabolites of this pathway in K562 (Extended Data Fig. 7b-e), MEL (Extended Data Fig. 7f-i), and primary mouse erythroid progenitor cells (Extended Data Fig. 7j), and confirmed the reduction of these metabolites. To test the role of choline in folate-depletion induced differentiation, we supplemented K562 with choline (500 μM) in high or low FA conditions. Supplementation with choline did not significantly alter proliferation (Extended Data Fig. 7k), or differentiation (Extended Data Fig. l) in both high and low FA conditions, suggesting that lack of choline is not a sufficient cause of folate-deprivation induced differentiation.

### Folate deprivation and 1C inhibition accelerate differentiation of primary murine erythroid progenitors

We next tested whether folate deprivation induces differentiation in primary erythroid progenitor cells (Fig. 4a). For this, we used an ex vivo murine erythroid progenitor differentiation system^29–31^ that allows controlled nutritional and pharmacological perturbations of 1C metabolism in primary progenitor cells with an established readout of differentiation. We isolated erythroid progenitors from fetal livers of E14.5 embryos and purified an enriched population of erythroid cells primarily composed of the erythroid-committed progenitor populations BFU-E (Burst Forming Unit-Erythroid) and CFU-Es (Colony Forming Unit – Erythroid)^32^. This ex vivo differentiation model of erythroid cells can be setup as two phases, expansion and differentiation, as defined by the growth factors present in the media. For addressing our biological question, we focused on culturing cells in the expansion phase to allow maintenance of undifferentiated progenitor cells for several days. The differentiation phase, achieved by removal of growth factors, is less suitable for our study because the strong differentiation drive is likely to mask any differentiation effect of 1C metabolism inhibition. The optimal commercial media commonly used for progenitor expansion (SFEM II) contains supraphysiological levels of folate. We therefore cultured the purified erythroid population in a custom folate-depleted RPMI-based expansion media that contains additional growth factors. Of note, erythroid progenitor cells cultured in this custom media supplemented with 2,000 nM FA proliferate successfully (Fig. 4b), but trend towards an early differentiation phenotype (Fig. 4c). Metabolite profiling of cells cultured in our custom expansion media supplemented with 0, 100, and 2,000 nM FA indicate the expected changes in 1C metabolites (Extended Data Fig. 8a-h). Expansion of primary progenitor cells occurs at a similar rate in the first two days of full or partial folate; however, by day 4 there is a significant proliferation defect in 100 nM and 0 nM FA compared to 2,000 nM FA (Fig. 4b). At day 4 folate-deprived progenitor cells increase expression of the differentiation marker Ter119 and decreased the progenitor cell marker TFRC (Fig. 4d, e), that together indicate advanced erythroid differentiation.

**Figure 4:**
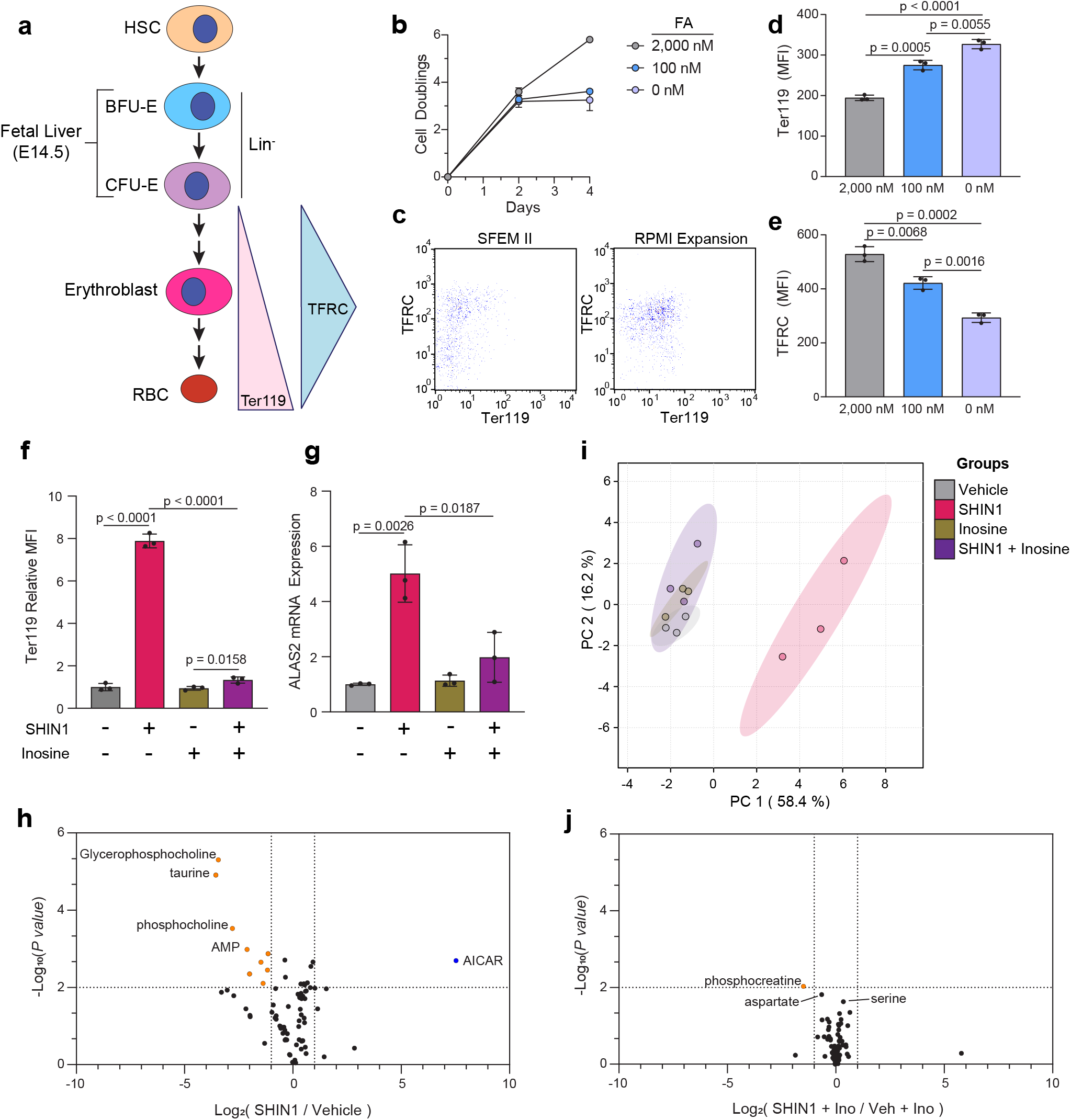
Folate deprivation induces differentiation in murine primary erythroid progenitor cells. **a,** Schematic representing murine fetal erythropoiesis from a population of Burst Forming Unit – Erythroid (BFU-E) and Colony Forming Unit-Erythroid (CFU-E) progenitor cells isolated from E14.5 fetal livers. Ter119 expression increases during erythroid differentiation. Transferrin receptor (TFRC) expression increases through the erythroblast stage, after which expression falls. RBC – red blood cell. **b,** Proliferation of murine primary erythroid progenitor cells in 2,000, 100, and 0 nM FA. **c,** Flow cytometry staining comparison between SFEM II media and custom RPMI-based expansion media supplemented with 2,000 nM FA. **d,** Ter119 surface expression and **e,** Transferrin Receptor (TFRC) expression at day 4 in 2,000, 100, and 0 nM FA measured by flow cytometry. **f,** Cell surface Ter119 levels on murine primary erythroid progenitor cells following 4 days treatment with vehicle, SHIN1 (1.25 μM), inosine (100 μM), or SHIN1 + inosine. **g,** RT-qPCR analysis of ALAS2 mRNA expression in murine primary erythroid progenitor cells treated for 4 days with SHIN1. **h,** PCA analysis of metabolite profiling data from murine primary erythroid progenitor cells cultured in expansion media with and without SHIN1 treatment and inosine supplementation. **i,** Metabolic changes of vehicle- or SHIN1-treated (1.25 μM, 2 days) murine primary erythroid progenitor cells cultured in SFEM II expansion media. **j,** Metabolic changes of vehicle- or SHIN1-treated (1.25 μM, 2 days) murine primary erythroid progenitor cells cultured in SFEM II expansion media supplemented with inosine (100 μM). Data shown are mean (± s.d.) of three biological replicates. All P values were calculated using an unpaired Student’s t-test.

To more closely investigate the effect of 1C metabolism inhibition on differentiation of early progenitor cells, we used the inhibitor SHIN1, because it allows culturing the cells in the commercial media SFEM II that maintains undifferentiated erythroid progenitor cells (Fig. 4c, Extended Data Figure 8i). SHIN1 treatment of primary progenitor cells induces differentiation that is rescued by inosine supplementation (Fig. 4f, g; Extended Data Figure 9a-c). Levels of the reduced folate 5-methyl THF are low in SHIN-1 treated progenitor cells, and this cannot be rescued by inosine supplementation (Extended Data Fig. 8j). Serine levels are high in SHIN1-treated cells, regardless of inosine supplementation, as expected from inhibition of SHMT – a serine metabolism enzyme (Extended Data Fig. 8k). This further supports the role of downstream nucleotides as mediators of differentiation as opposed to signaling induced by direct sensing of the depleted folate levels. Inosine supplementation significantly increased hypoxanthine levels, in agreement with the metabolic fate of inosine as a substrate for salvage purine synthesis (Extended Data Fig. 8l). As in folate-deficient cells, SHIN1-treated cells show AICAR accumulation, and reduced levels of glycerophosphocholine and phosphocholine (Fig. 4i, j). Supplementation with inosine rescues nearly all metabolic changes associated with SHIN1 treatment (Fig. 4h), which further suggests the role of purine synthesis impairment in regulating differentiation. As shown in K562 and MEL cells, SHIN1 treatment had no effect on IMP levels, and only a modest effect on NTP levels. The effect of SHIN1 on ATP and GTP was rescued with inosine supplementation (Extended Data Fig. 8m, n). Importantly, SHIN1 treatment induces differentiation of primary erythroid progenitor cells, and this can be reversed by inosine supplementation (Fig.4f, g; Extended Data Figure 9a-c). Taken together, folate depletion and 1C inhibition induce differentiation in primary early erythroid progenitors. This induction of differentiation can occur even in the presence of growth factors and is blocked by supplementation with exogenous nucleosides, which suggests that purine levels are sensed, and their depletion activates premature differentiation.

## Discussion

Deprivation of the essential vitamin folate results in megaloblastic anemia in children and adults^33,34^. This disease remains a significant health incumbrance in underdeveloped countries^35,36^, and in underserved populations, where nutritional shortage of folate still prevails even in our era of folate fortification and common supplementation^9,37^. Megaloblastic anemia is characterized by the appearance of big, hemoglobin-filled, erythrocytes^38,39^. Early studies have suggested folate-deficient anemia is a result of DNA damage induced apoptosis^6,7^. It has been thought that the absence of folate decreases thymidylate synthesis and subsequently, the ratio of dTTP:dUTP. As a result, uracil is misincorporated into DNA, induces DNA damage, and leads to increased erythroid apoptosis, which manifests as anemia. However, although several studies have called into question this mechanism of folate-deficient anemia because of lack of molecular evidence for both DNA damage and apoptosis of erythroid progenitors^40,41^, no alternative hypothesis has yet emerged.

Our data indicate that folate deprivation induces a metabolic stress that results in aberrant differentiation of erythroid cells. Premature differentiation of progenitor cells can result in insufficient number of cell division cycles that normally proceed differentiation, and reduced cell numbers of cells of that lineage. Our results suggest an etiology to the low cell count of erythroid cells in megaloblastic anemia, and to the unique manifestation of this disease – large, heme-filled, cells. We describe here a folate deprivation-induced increase in heme metabolism that is regulated transcriptionally. We have not yet identified the upstream signaling cascade that is driving the transcriptional response to folate deprivation, but we postulate that the drop in cellular folate is being sensed, either directly or through purine levels, leading to signal transduction and activation of a differentiation program. Our data rule out the involvement of two major signaling pathways that respond to changes in purine levels – the mTORC1 and AMPK pathways – and we postulate the existence of a full signaling cascade that is bridging between the drop in folate availability and activation of the differentiation program in erythroid cells.

By providing cells in culture with folate concentration that might mimic the mild shortage in folate availability as in nutritional folate deprivation, and by comprehensive profiling of the metabolic consequences of this mild folate deprivation in erythroid cells, we identified premature, aberrant differentiation as a possible etiology for nutritional megaloblastic anemia.

## Methods

### Cell lines

All cell lines were tested monthly and found to be negative for mycoplasma infection. The sources of the cell lines are as follows: K562, HEL, RAMOS, P12-Ichikawa, MOLM13, NALM6, NOMO, EOL-1: D.M. Sabatini, MIT; MEL: M.D. Fleming, BCH. All cells were cultured at 37 °C with 5% CO2.

### Cell Culture Experiments

Cell lines were maintained in folate-free RPMI (Sigma-Aldrich, R1145) supplemented with 10% dialyzed FBS, Sodium Bicarbonate (Sigma-Aldrich, S8761), L-glutamine (Sigma-Aldrich, G7513), and the indicated concentration of FA (Sigma-Aldrich, F8758). The reagents used in cell culture experiments are as follows: Ferric ammonium citrate (Sigma-Aldrich, F5879), iron-bound transferrin (Sigma-Aldrich, T4132), deferoxamine (Sigma-Aldrich, D9533), inosine (Sigma-Aldrich, I4125), AICAR (Cayman Chemical, 10010241), SHIN1 (MedChem Express, HY-112066), Rapamycin (EMD Millipore, 553210), TORIN1 (Cayman Chemical, 10997), GSK621 (Sigma-Aldrich, SML2003), choline (Sigma-Aldrich, C7527), [2-^13^C]serine (Cambridge Isotope Laboratories, CLM-2013-PK), [2-^13^C]glycine (Cambridge Isotope Laboratories, CLM-136-PK). All cell culture experiments were seeded with a starting cell density of 100,000 cells per mL and cultured for 2 days in the specified culture media. After 2 days, cells were counted, and reseeded at 100,000 cells per mL in the specified condition with fresh media. Cell proliferation studies continued for 6 days and cell doubling calculations were calculated using the equation:

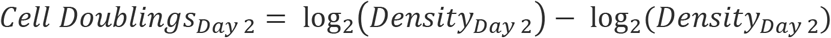

Cumulative doublings were calculated by summing the cell doublings between day 0 and day 6.

### Ex vivo expansion

Isolation and purification of lineage depletion, erythroid progenitor cells from fetal livers of E14.5 mice was performed as previously described^24^. In brief, fetal livers are homogenized in PNEG buffer (PBS, 20 μM EDTA, 2% neonatal calf serum, 10 mM glucose) and depleted of red blood cells by red blood cell lysis. Lineage depletion is achieved through magnetic pull down with streptavidin beads and the following biotin conjugated antibodies: mouse lineage depletion panel [CD3e, CD45R, Ly-6G, Ly-6C, CD11b, Ter119] (BD Biosciences, 559971), CD16/32, clone 93 (E-Biosciences, 13-0161-82), CD41, clone eBioMWReg30 (E-Biosciences, 13-0411-82), Ter119 (E-Biosciences, 13-5921-82), Ly-6A/E, clone D7 (E-Biosciences, 13-5981-82). The clarified supernatant that contains CFU-E and BFU-E erythroid progenitors is retrieved, washed, and stored on ice until cell culture media is prepared. Progenitor cells were cultured in either StemSpan SFEM II Expansion media (StemCell Technologies, 09605), or our custom, folate-free RPMI based expansion media. Both SFEM II and our custom expansion media were supplemented with 100 ng/mL stem cell factor (Peprotech, 250-03), 40 ng/mL insulin-like growth factor (Peprotech, 250-19), 100 mM dexamethasone (Sigma-Aldrich, D9184), and 2 U/mL erythropoietin (Peprotech, 100-64). Our custom RPMI-based expansion media was additionally supplemented with 10 μg/mL insulin (Sigma-Aldrich, I9278) and 250 μg/mL saturated human holo-transferrin (Sigma-Aldrich, T4132). Progenitor cells were seeded in the respective media conditions at a density of 50,000 cells per mL. Cell proliferation was assessed by cell counts at day 2 and day 4 of expansion.

### Metabolite Profiling by mass spectrometry

#### Polar metabolite detection

One million cells from culture were collected via centrifugation, washed with 0.9% NaCl, and resuspended in extraction buffer (80% Methanol, 25 mM Ammonium Acetate and 2.5 mM Na-Ascorbate prepared in LC-MS water, supplemented with isotopically-labelled amino acid standards [Cambridge Isotope Laboratories, MSK-A2-1.2], aminopterin, and reduced glutathione standard [Cambridge Isotope Laboratories, CNLM-6245-10]). Samples were vortexed for 10 sec, then centrifuged for 10 minutes at 18,000 g to pellet cell debris. The supernatant was divided into two tubes and dried on ice using a liquid nitrogen dryer. One tube of dried sample was saved for folates metabolite detection (see below), and the second was used for polar metabolite detection. Dried samples were resuspended in 25 μL water and 2 μL was injected into a ZIC-pHILIC 150 × 2.1 mm (5 μm particle size) column (EMD Millipore). operated on a Vanquish™ Flex UHPLC Systems (Thermo Fisher Scientific, San Jose, CA, USA). Chromatographic separation and MS data acquisition was performed as previously described^42^.

#### Folate metabolite detection

Dried samples were prepared for folate detection similarly to that previously described^43^. The mass spectrometer was operated in full-scan, positive ionization mode using three narrow-range scans: 438–450 m/z; 452–462 m/z; and 470–478 m/z, with the resolution set at 70,000, the AGC target at 10e6, and the maximum injection time of 150 ms. HESI settings were: sheath gas flow rate: 40; Aux gas flow rate: 10; Sweep gas: 0; Spray voltage: 2.8 (neg) 3.5 (pos); Capillary temperature 300; S-lens RF level 50; Aux gas heater temp: 350. Levels of folates were normalized to aminopterin as an internal standard and to polar metabolites.

#### Porphyrin metabolite detection

Porphyrin extraction was based on published protocol^44^, with modifications. In brief, one million cells from culture were collected via centrifugation, washed with 0.9% NaCl, and resuspended in 150 μl of porphyrin extraction buffer (1:4 ratio of 1.7 M HCl:ACN, 1μM deuteroporphyrin IX (Frontier Scientific, D510-9)) and 0.5 μM isotopically labeled amino acids (Cambridge Isotopes, MSK-A2-1.2)). Samples were vigorously shaken for 20 min at 16°C in a thermomixer (Eppendorf), sonicated for 10 cycles at 4°C with 30 sec on and 30 sec off, then incubated at 4°C for 10 min. Following incubation on ice, samples were centrifuged for 10 minutes at 18,000g to pellet cell debris. The supernatant was collected and 40.5 μl super-saturated MgSO4 and 12 μl 5 M NaCl were added. Samples were vortexed for 30 sec and further shaken for 10 min at 16 °C in a thermomixer. Finally, a 10 min 10,000 rpm centrifugation was used to separate the organic layer (upper) from the aqueous layer (lower). The upper organic layer was collected and 5 uL was injected onto a 2.6 μm, 150 × 3 mm C18 column (Phenomenex, 00F-4462-Y0) equipped with a 3.0 mm safe-guard column (Phenomenex, AJ0-8775). Column compartment was heated to 45 °C. Porphyrins were separated with a chromatographic gradient at a flow rate of 0.800 ml min-1 as follows: 0–2 min: 5% B; 2-19min: linear gradient from 5% to 95% B; 19-21min: 95% B; 21.1-23min: return to 5% B. The mass spectrometer was operated in full-scan, positive ionization mode using a narrow-range scan: 450-700m/z, with an additional tSIM scan for hemin (616.1767 m/z), CoproP (655.2762 m/z), and PPIX (563.2653 m/z) with the resolution set at 70,000, the AGC target at 1e6, and the maximum injection time of 50 ms. HESI settings were: sheath gas flow rate: 40; Aux gas flow rate: 10; Sweep gas: 0; Spray voltage: 2.8 (neg) 3.5 (pos); Capillary temperature 300; S-lens RF level 55; Aux gas heater temp: 350.

### Metabolomics Data Analysis

Polar metabolites, folates and porphyrins were relatively quantified while referencing an in-house library of chemical standards and using TraceFinder 4.1 (Thermo Fisher Scientific, Waltham, MA, USA), with a 5 ppm mass tolerance. Pooled samples and fractional dilutions were prepared as quality controls and injected at the beginning and end of each run. Pooled samples were interspersed throughout the run to control for technical drift in signal quality as well as for coefficient of variability (CV) determination for each metabolite. Data normalizations were performed in two steps; 1. Integrated peak area signal from internal standards added to extraction buffers were mean-centered (for every standard, peak area was divided by the mean peak area of the set) and averaged across samples; samples were divided by the resulting factor, thus normalizing for any technical variability due to MS-signal fluctuation or pipetting and sample injection errors (usually within 10% variability). 2. Normalization for biological material was based on detected polar metabolites as follows: CV values (based on pooled sample re-injections) and coefficient of determination (RSQ) (based on linear dilutions of pooled sample) were calculated per metabolite, metabolites with <30% CV and >0.95 RSQ were mean-centered and averaged across samples. Metabolite peak areas were then divided by the resulting factor (biological normalizer), thus accounting for any global shift in metabolite amounts due to differences in biological material.

### Serine and Glycine tracing

Tracing experiments were conducted in folate, B12, and amino acid-free RPMI (US Bio, R9010-06) supplemented with 10% dialyzed FBS, B12 (Sigma Aldrich, V6629), amino acids (to the standard RPMI concentration, see table below for product information). Prior to tracing, cells were cultured as described above in amino acid-free RPMI supplemented with all unlabeled amino acids. [2-^13^C] serine and [2-^13^C]glycine tracing was performed in K562 cells following 8 days culture in 2,000 nM or 100 nM FA. At day 8, cells were washed and placed in amino acid-free RPMI at the indicated FA concentration containing all amino acids minus serine and glycine. As indicated, cells were supplemented with: [unlabeled]serine + [unlabeled]glycine, [2-^13^C]serine + [unlabeled]glycine, or [unlabeled]serine + [2-^13^C]glycine. Total glycine and serine concentrations in all media conditions were 10 mg/L and 30 mg/L, respectively. Following 24hr culture in labeled or unlabeled conditions, cells were quickly washed in 0.9% NaCl and extracted in either polar or porphyrin extraction buffer. Relative quantification of polar metabolites was performed with TraceFinder 4.1 as described above. To identify metabolites with expected ^13^C labeling, the mass of the extra proton (m = 1.00727) was added to the expected *m/z* for each potential carbon that could be labeled. Relative quantification of this list of compounds allowed the calculation of percent labeling for each carbon (m+1, m+2, m+3, etc.). Subtraction of the natural abundance of 13C was performed using the R package, IsoCorrectoR^45^. Corrected abundances were presented as the change in percent labeling between experimental conditions.

### Immunoblot

Cell lysis was performed with RIPA buffer (Santa Cruz, sc-24948) following manufacturer’s protocol. Protein concentration was measured using BCA assay (Thermo, 23227). Protein samples were prepared in 2X Sample buffer (Thermo, LC2676) so that 20 ug protein lysate per sample could be used for SDS-PAGE. Immunoblotting was performed using the following primary antibodies: hemoglobin A1 (Abcam, ab92492; 1:1,000), hemoglobin, fetal (Abcam, ab137096; 1:1,000), p-ULK1-S555 (CST, 5869T; 1:1,000), ULK1 (CST, 8054S; 1:1,000), p-ACC-S79 (CST, 3661P; 1:1,000), ACC (CST, 3676P; 1:1,000), vinculin (CST, 13901; 1:2,500), p-S6-T389 (CST, 9205; 1:1,000), S6 (CST, 9202; 1:1,000), AMPKa (CST, 5832; 1:1,000), SHMT2 (CST, 33443; 1:1,000). HRP-conjugated anti-rabbit (Jackson ImmunoResearch, 111-035-144; 1:10,000) and anti-mouse (Jackson ImmunoResearch, 115-035-166; 1:10,000) secondary antibodies were used.

### Flow Cytometry

Flow cytometry was performed on a BD FACSCelesta. In brief, cells were washed with FACs buffer (PBS +2% FBS) and incubated for 20 minutes in the presence of the indicated antibodies. Cells were washed in FACS buffer and viability was assessed using 1 μg/mL DAPI (Sigma-Aldrich, D9542). The following flow cytometry antibodies were used: Ter119 (1:200; BioLegend, 116206), Glycophorin A (1:200; BioLegend, 349104), Transferrin Receptor (mouse) (1:200; BioLegend, 113819), Transferrin receptor (human) (1:200; BioLegend, 334108).

### Gene Expression Analysis (RNA, cDNA, qPCR, analysis)

RNA extraction from cell pellets were performed using RNAzol (Fischer, NC0477546). The reverse transcriptase reaction was performed using between 100 ng and 1 μg RNA per sample (NEB, E3010). Quantitative PCR (qPCR) reactions were performed with SYBR green (CWBiotech, CW2621) on the QuantStudio 7 Pro (Applied Biosystems) qPCR instrument. The reference gene for all qPCR reactions were beta actin.

Primers used for qPCR:

hActin

Forward – CAACCGCGAGAAGATGACCC
Reverse – AGGCGTACAGGGATAGCACA
mActin

Forward – GGCTGTATTCCCCTCCATCG
Reverse - CCAGTTGGTAACAATGCCATGT
hALAS2:

Forward – CACCACAGCCCTCAGATGAT
Reverse – CAAAGTGTACAGGACGGCGA
mALAS2:

Forward – GGGCTAAGAGCCATTGTCCT
Reverse – ATGGCTTCGGGTGGTTGAAT
hALAS1:

Forward – AGATCTGACCCCTCAGTCCC
Reverse – TCCACGAAGGTGATTGCTCC
ALAD:

Forward – GGCTACTTCCACCCACTACTT
Reverse – ATGGAAGTGGAGATGGTGGGA
HMBS:

Forward – GCAGGAGTTCAGTGCCATCATC
Reverse – AGCACACCCACCAGATCCAAG
UROS:

Forward – CAGTTGCACACCCAGGAATC
Reverse – ATGTGAGGCCAGAGGGACTAA
UROD:

Forward – GCTGACCATGGAAGCGAATG
Reverse – TGCACCAAACGGGAGTGTAG
CPOX:

Forward – TGGGCGTGAGCTCTGTTATC
Reverse – CACCAAACCACCACTGCTTG
PPOX:

Forward – GAACAGGTTCCTCTACGTGGG
Reverse – GAGACGCCACCTCAGGTCC
FECH:

Forward – AAAGGGCTTTGTGAGAGGGG
Reverse – CAGGGCCTTAGAGAACAATGGA
MTHFS:

Forward – GAGTGATTGCCCACAGTGAGT
Reverse – TTTGCCTCGTTGGAAAATGT
SHMT1:

Forward – GAGCAGTTTTGGAGGCCCTA
Reverse – CTCGCTTCTGACAGAGGGTC
SHMT2:

Forward – CCTAGACCAGAGTTGGTGGC
Reverse – TGTTTGCTTCCCCAGTCTGA
MTHFD1:

Forward – GTGCACTCACTGGGCAGAAG
Reverse – TCAGCTTTGTGTTGAGCTTCG
MTHFD2:

Forward – GGTCTATGGCTGCGACTTCTCT
Reverse – TTGATCTGCTGGGCCAGTTTC
SLC19A1:

Forward – GAGAGCTTCATCACCCCCTA
Reverse – TGTGCATCTCCCAAAGTGTG
TYMS:

Forward – GGTGTGCCTTTCAACATC
Reverse – GGCCAGGTAGGAGTACGACA

### Mitochondrial Analysis

Mitochondrial mass was assessed using the mitochondrial permeable, potential independent dye Mitospy (BioLegend, 424805). Mitochondrial potential was assessed using the potential dependent dye, Mitotracker deep red (Invitrogen, M22426). Cells were incubated at 37°C for 30 minutes in the presence of dye and then prepared for flow cytometry as described above. The Seahorse XFe96 Analyzer was used to assess oxygen consumption rate (OCR) in K562. Prior to OCR measurement, cells were washed and placed in a custom, folate-free RPMI media lacking glutamine, pyruvate, glucose, phenol red, and sodium bicarbonate (Gibco). The media was supplemented with pyruvate, glucose, and l-glutamine immediately prior to experiment. K562 were adhered to the 96 well plate using Cell-Tak then incubated at 37°C for 1 hour in an incubator without CO2. The standard mitochondrial stress test was performed with injection of FCCP (2 μM; Sigma-Aldrich, C2920), oligomycin (1 μM; Sigma-Aldrich, 75351), rotenone (500 nM; Sigma-Aldrich, R8875) and Antimycin A (500 nM; Sigma-Aldrich, A8674). Analysis was performed using Wave software (Agilent Technologies).

### Statistical analysis and software

Analysis and visualization of metabolomics data was generated with MetaboAnalyst 5.0 (www.metaboanalyst.ca). Statistical analysis was performed using GraphPad Prism 9. For pairwise comparisons, two-tailed Student’s t-tests were used. For multiple comparisons, one-way ANOVA test with Tukey’s post-hoc test was used. All error bars represent standard deviation (S.D.).

## Data Availability

Original western blot scans are attached as supplementary figures. Raw qPCR data are attached as source data, original FACS plots are attached as supplementary files or added to Extended Data Figures. Raw metabolomics data are uploaded to the NIH Common Fund’s National Metabolomics Data Repository (NMDR) and are available at the Metabolomics Workbench Metabolite Database (https://www.metabolomicsworkbench.org/databases/metabolitedatabase.php).

## Amino Acid Table (Sigma-Aldrich)References

**Amino Acid Table (Sigma-Aldrich)**
|  |  |
| --- | --- |
| Glycine | G8790-100G |
| L-Arginine | A8094-25G |
| L-Asparagine | A4159-25G |
| L-Aspartic acid | A7219-100G |
| L-Cystine 2HCl | C6727-25G |
| L-Glutamic Acid | G8415-100G |
| L-Glutamine | G8540-100G |
| L-Histidine | H6034-25G |
| L-Hydroxyproline | H5534-25G |
| L-Isoleucine | I7403-25G |
| L-Leucine | L8912-25G |
| L-Lysine hydrochloride | L8662-25G |
| L-Methionine | M5308-25G |
| L-Phenylalanine | P5482-25G |
| L-Proline | P5607-25G |
| L-Serine | S4311-25G |
| L-Threonine | T8441-25G |
| L-Tryptophan | T8941-25G |
| L-Tyrosine disodium salt dihydrate | T1145-25G |
| L-Valine | V0513-25G |

## Acknowledgements

We thank all members of the Kanarek lab for their advice and help; We thank Dr. Jose M. Orozco for the AMPK KO plasmids; We thank Drs David M. Sabatini and Raul Mostoslavsky, for reading and editing the manuscript. This work was supported by funding from The Charles H. Hood Foundation (N.K.); Gabrielle’s Angels Foundation for Cancer Research #135 (N.K.); The Smith Family Awards Program for Excellence in Biomedical Research (N.K.); Research and Recruitment Funding by BCH (N.K.); Landry Cancer Biology Consortium (A.M.); N.K. is a Pew Biomedical Scholar.

## Author Information

^1^Department of Pathology, Boston Children’s Hospital, Boston, MA, 02115, USA

^2^Graduate Program in Biological and Biomedical Sciences, Harvard Medical School, Boston, MA, 02115, USA

^3^Harvard Medical School, Boston, MA, 02115, USA

^4^Harvard MD-PhD Program, Harvard Medical School, Boston, MA, 02115, USA

^5^Koch Institute for Integrative Cancer Research, Massachusetts Institute of Technology, Cambridge, MA 02142, USA.

^6^Division of Hematology/Oncology, Boston Children’s Hospital, Boston, MA, 02115, USA

^7^Department of Pediatric Oncology, Dana-Farber Cancer Institute, Boston, MA, 02115, USA

^8^Broad Institute of Harvard and MIT, Cambridge, MA, USA.

## Contributions

A.G.M and N.K formulated the research plan and interpreted experimental results; A.G.M designed and performed all experiments with assistance from N.K.P., B.P., A.W., P.W., and A.J.C.; B.P. operated the metabolite-profiling instruments and advised and helped planning experiments for metabolites detection by LC-MS; A.J.C. uploaded raw metabolomics data on a public repository; L.H., N.H., O.K. and T.B. set up mouse breeding and performed cell purifications of murine fetal liver primary hematopoietic stem cells; S.D., M.D.F. and H.L. provided guidance and reagents in regards to heme and iron metabolism, and to anemia, metabolic diseases, and hematopoiesis; B.P. helped with figure design and layout. A.G.M and N.K wrote the manuscript, and all the authors edited it.

## Ethics Declarations

Animal work performed as part of this study was done in accordance with all ethical guidelines and regulations. No human samples or participants were involved in the study.

