## Extended Data Figures for "Folate depletion induces erythroid differentiation through perturbation of de novo purine synthesis"

**Abstract**

All dividing cells require the essential vitamin folate. Hematopoietic cells harbor a unique sensitivity to folate deprivation, as implied by the development of folate-deficient anemia and the utility of anti-folate chemotherapy in blood cancer. To study this metabolic sensitivity, we applied mild folate depletion to

### Extended Data Figure 1

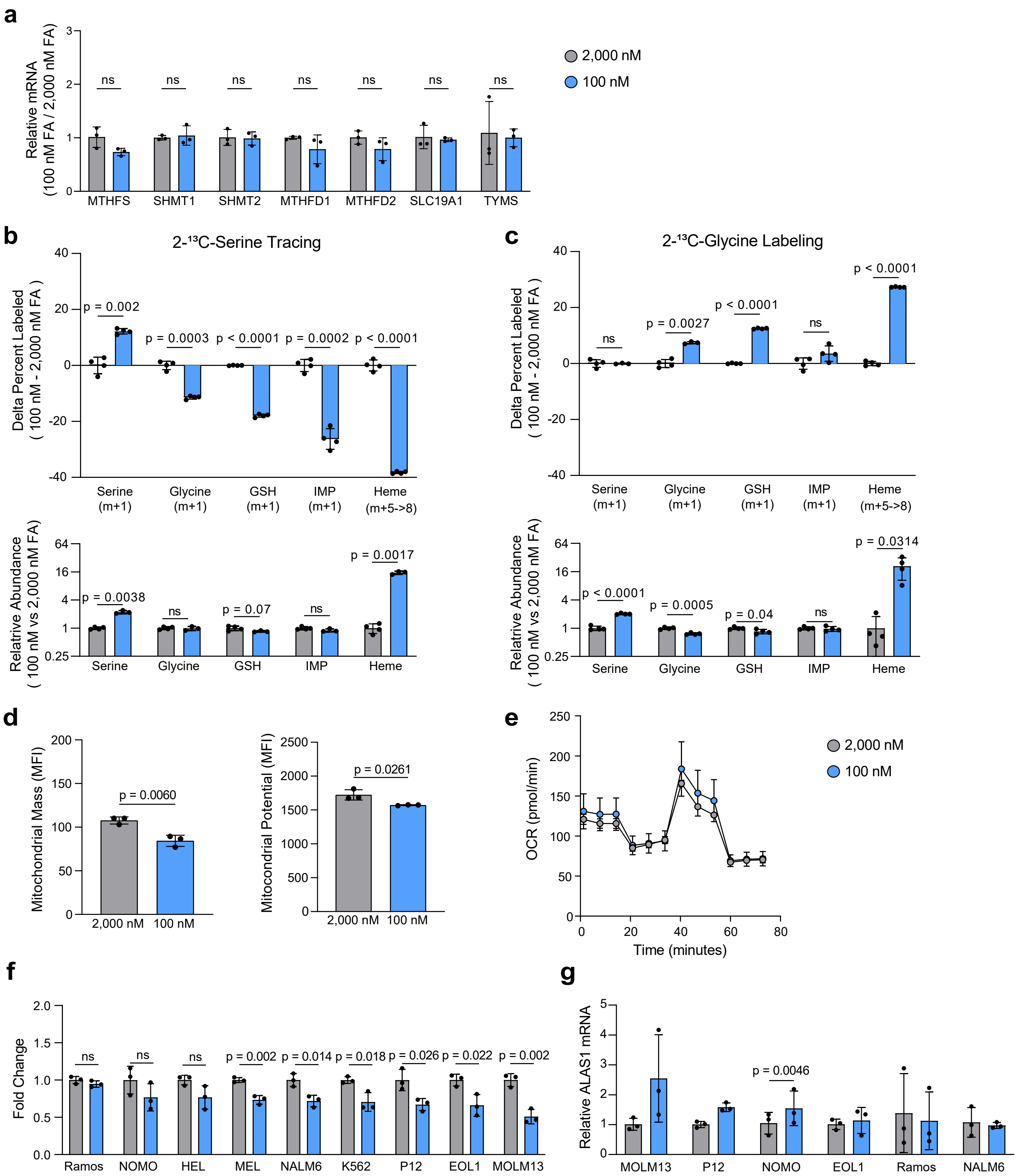

**Extended Data Figure 1.**

**a,** RT-qPCR analysis of 7 genes related to 1-carbon (1C) metabolism following 6 days culture in 2,000 and 100 nM FA. **b, c,** Difference in $^{13}C$-labeling (top) or total levels (bottom) in serine, glycine, glutathione (GSH), inosine monophosphate (IMP), and Heme from (**b**) 2-$^{13}C$-Glycine and (**c**) 2-$^{13}C$-Serine following 24hr isotope labeling in K562 cells cultured in 2,000 or 100 nM FA. **d,** Flow cytometry measurement of mitochondrial mass and membrane potential as quantified by mean fluorescence intensity (MFI) of Mitospy (mass) and Mitotracker (membrane potential). Data shown for K562 in 2,000 and 100 nM FA at day 6. **e,** Oxygen consumption rate (OCR) from the Seahorse mitochondrial stress test in K562 at day 6 in 2,000 and 100 nM FA. **f,** Relative proliferation rate of nine leukemia cell lines over 6 days in 2,000 and 100 nM FA. **g,** RT-qPCR analysis of ALAS1 mRNA expression in non-erythroid cell lines. Data shown are mean (± s.d.) of three biological replicates. All P values were calculated using an unpaired Student's t-test.

### Extended Data Figure 2

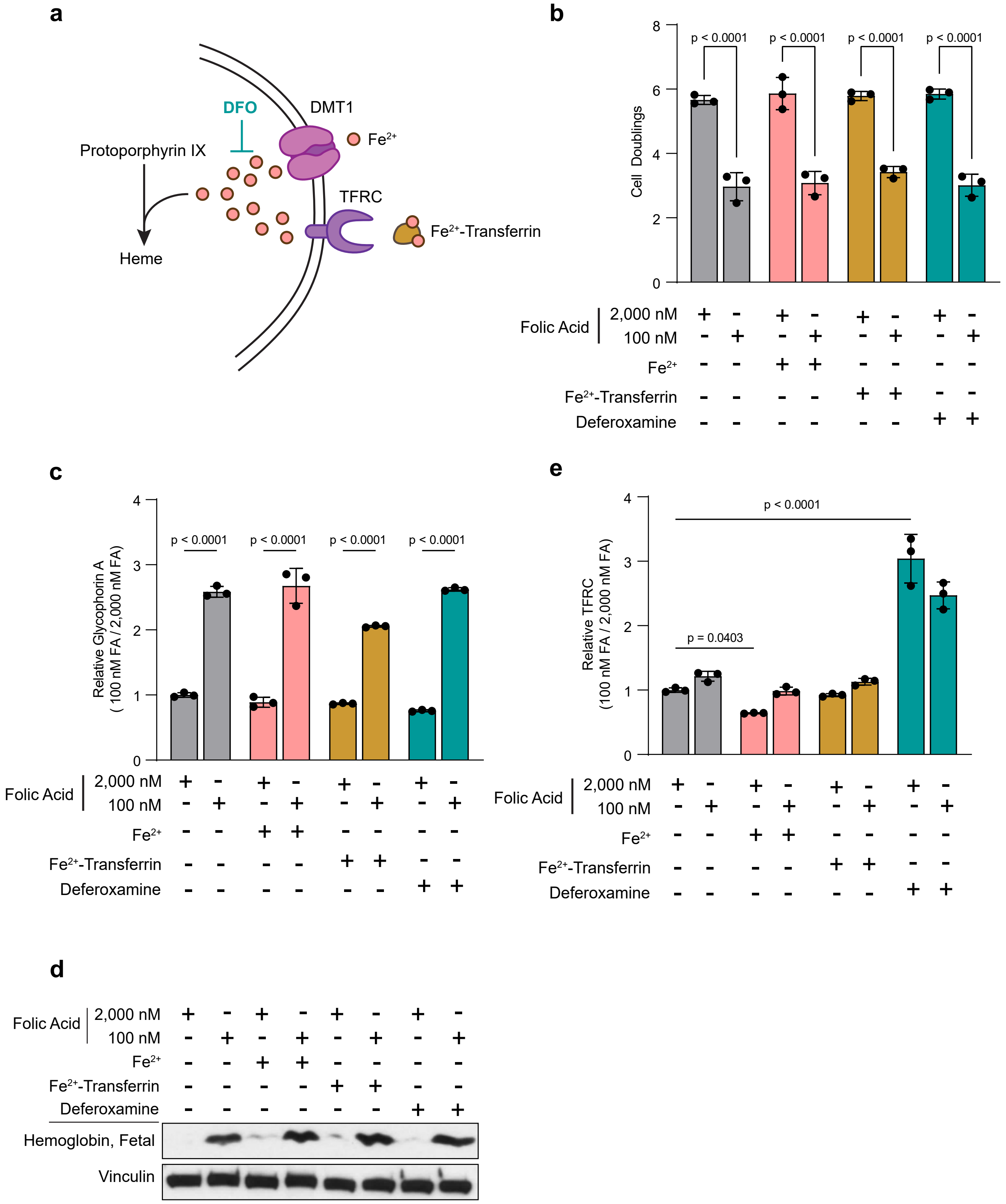

**Extended Data Figure 2.**

**a,** Schematic depicting the role of iron intake and utilization in heme biosynthesis. **b-e,** Cell proliferation (**b**), glycophorin A expression (**c**), hemoglobin expression (**d**), and transferrin receptor (TFRC) expression (**e**) in K562 following 6 days culture in 2,000 and 100 nM FA media supplemented with free iron (Ammonium Ferric Citrate), iron-bound transferrin, and the iron chelator, deferoxamine (DFO). Data shown are mean (± s.d.) of three biological replicates. All P values were calculated using an unpaired Student's t-test.

#### Extended Data Figure 3

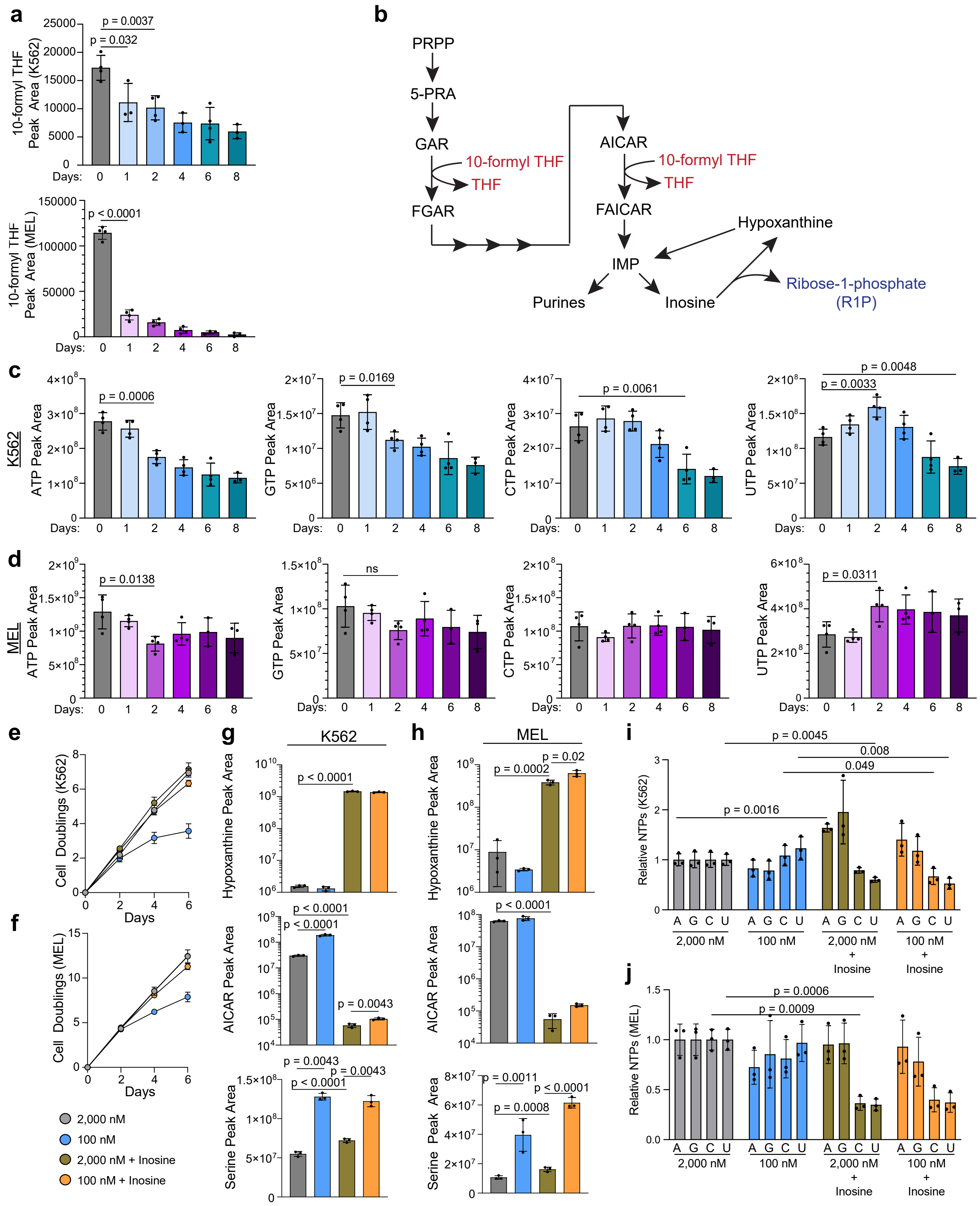

**Extended Data Figure 3.**

**a,** 10-formyl THF levels in K562 (top) and MEL (bottom) through day 8 in 100 nM FA as measured by LC-MS. **b,** Simplified schematic depicting the purine synthesis and salvage pathways. Highlighted are the folate co-factors necessary for two of the reactions in this pathway. **c, d,** Nucleotide triphosphate levels in K562 (**c**) and MEL (**d**) measured by LC-MS at day 8 in 100 nM FA. **e, f,** Proliferation of K562 (**e**) and MEL (**f**) over 6 days in 2,000 and 100 nM FA alone, or with inosine supplementation. **g, h,** Hypoxanthine, AICAR, and Serine levels in K562 (**g**) and MEL (**h**) following 2 days in 2,000 and 100 nM FA with or without inosine supplementation. **i, j,** Relative levels of nucleotide triphosphate after 2 days in 2,000 and 100 nM FA, with or without inosine supplementation in K562 (**i**) and MEL (**j**). Data shown are mean (± s.d.) of three biological replicates. All P values were calculated using an unpaired Student's t-test.

### Extended Data Figure 4

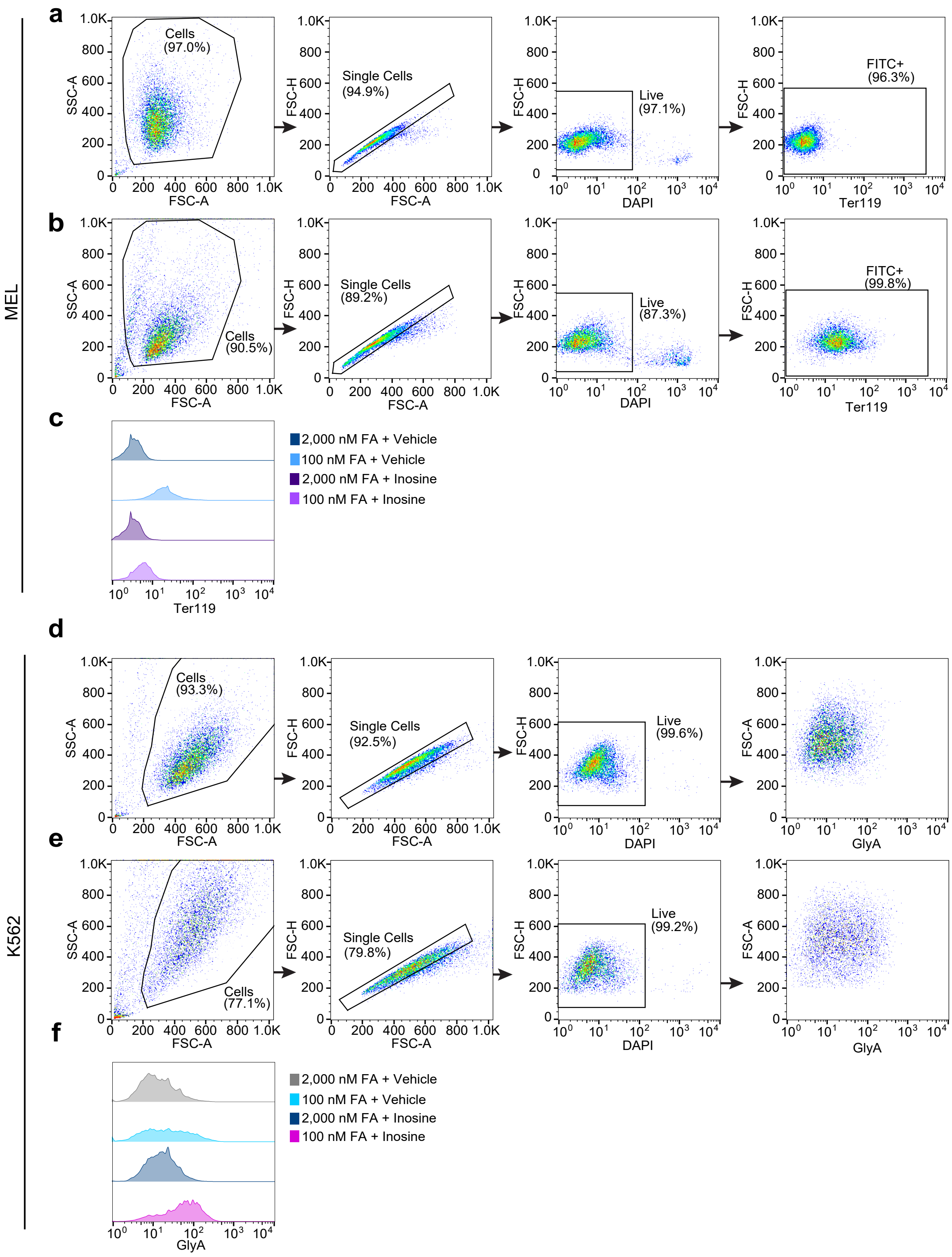

**Extended Data Figure 4.**

**a, b,** Representative flow cytometry populations showing MEL cells in 2,000 nM FA (**a**) or 100 nM FA (**b**) for 6 days. **c,** Representative histograms of Ter119 expression from MEL cells cultured for 6 days in 2,000 nM FA, 100 nM FA, 2,000 nM FA + Inosine, or 100 nM FA + Inosine. **d, e,** Representative flow cytometry populations showing K562 cells in 2,000 nM FA (**d**) or 100 nM FA (**e**) for 6 days. **f,** Representative histograms of GlycophorinA expression from K562 cells cultured for 6 days in 2,000 nM FA, 100 nM FA, 2,000 nM FA + Inosine, or 100 nM FA + Inosine.

#### Extended Data Figure 5

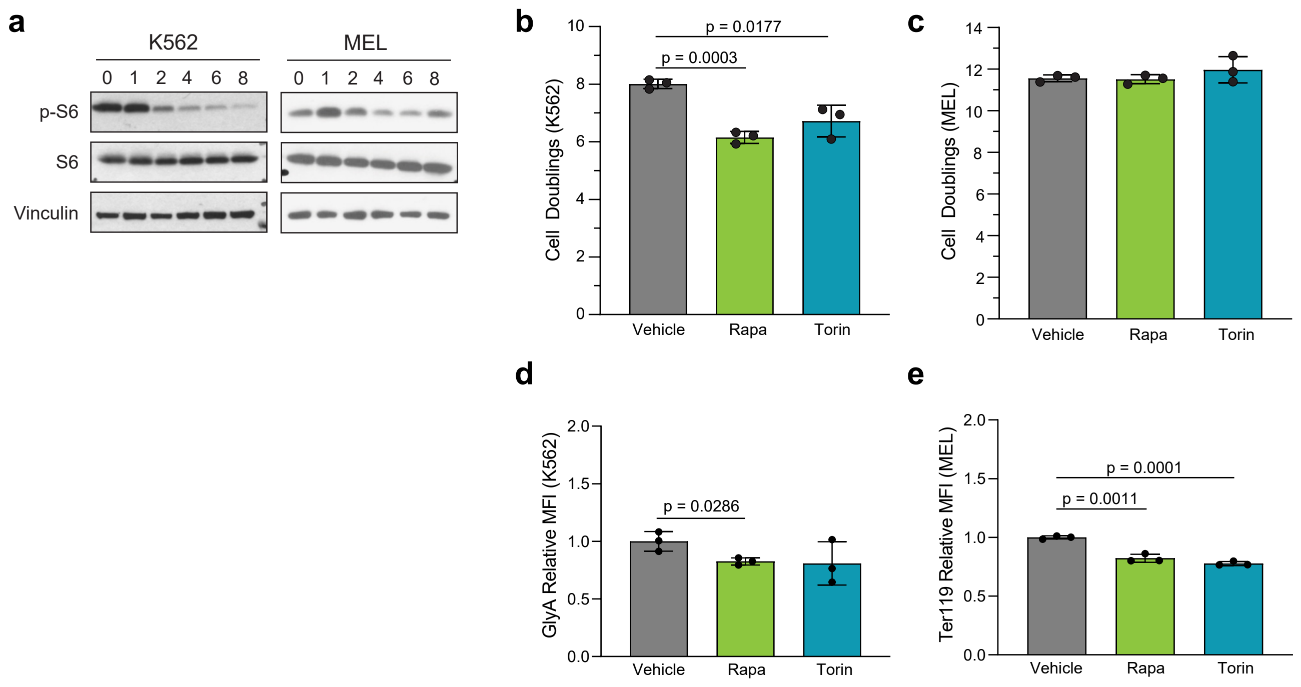

**Extended Data Figure 5.**

**a,** Western blot analysis of phospho-S6 and total S6 levels in K562 and MEL over 8 days in 100 nM FA. Vinculin is a loading control. **b, c,** Cell proliferation of K562 (**b**) and MEL (**c**) at day 6 in vehicle, Rapamycin (100 µM), and Torin1 (5 nM) treatments. Media and drug treatments were refreshed every 2 days. **d,** Cell surface Glycophorin A levels on vehicle-, Rapamycin-, and Torin1-treated K562 (for 6 days). **e,** Cell surface Ter119 levels on vehicle-, Rapamycin-, and Torin1-treated MEL (for 6 days). Data shown are mean (± s.d.) of three biological replicates. All P values were calculated using an unpaired Student's t-test.

### Extended Data Figure 6

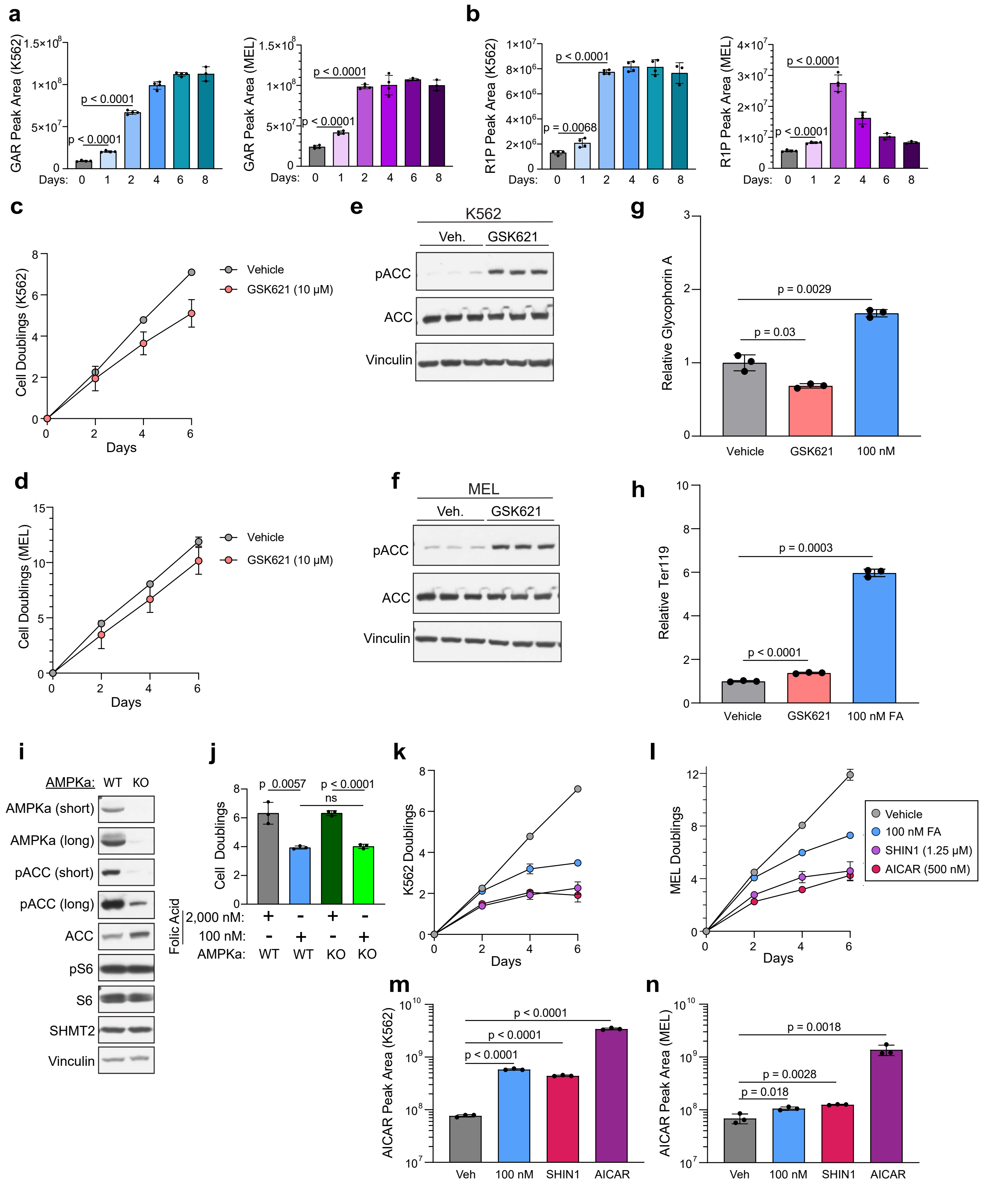

**Extended Data Figure 6.**

**a,** GAR levels in K562 (left) and MEL (right) over 8 days in 100 nM FA measured by LC-MS. **b,** Ribose-1-phosphate (R1P) levels in K562 (left) and MEL (right) over 8 days in 100 nM FA measured by LC-MS. **c, d,** Proliferation of vehicle- and GSK621-treated (10 µM) K562 (**c**) and MEL (**d**) (for 6 days). **e, f,** Western blot analysis of phospho-ACC/ACC, with and without GSK621 treatment in K562 (**e**) and MEL (**f**). Vinculin is a loading control. **g, h,** Cell surface levels of Glycophorin A (GlyA) (**g**) and Ter119 (**h**) measured by flow cytometry of K562 (**g**) and MEL (**h**) cultured in 2,000 nM FA + vehicle, 2,000 nM FA + GSK621 (10 µM), and 100 nM FA. **i,** Western blot analysis of AMPKa1/a2 WT and DKO K562. Vinculin is a loading control. **j,** Proliferation of AMPKa1/a2 WT and DKO K562 in 2,000 and 100 nM FA over 6 days. **k, l,** Proliferation of K562 (**k**) and MEL (**l**) over 6 days in 2,000 nM FA + vehicle, 100 nM FA, 2,000 nM FA + SHIN1 (1.25 µM), and 2,000 nM FA + AICAR (500 mM). **m, n,** AICAR levels in K562 (**m**) and MEL (**n**) following 2 days in the treatments listed above as measured by LC-MS. Data shown are mean (± s.d.) of three biological replicates. All P values were calculated using an unpaired Student's t-test.

### Extended Data Figure 7

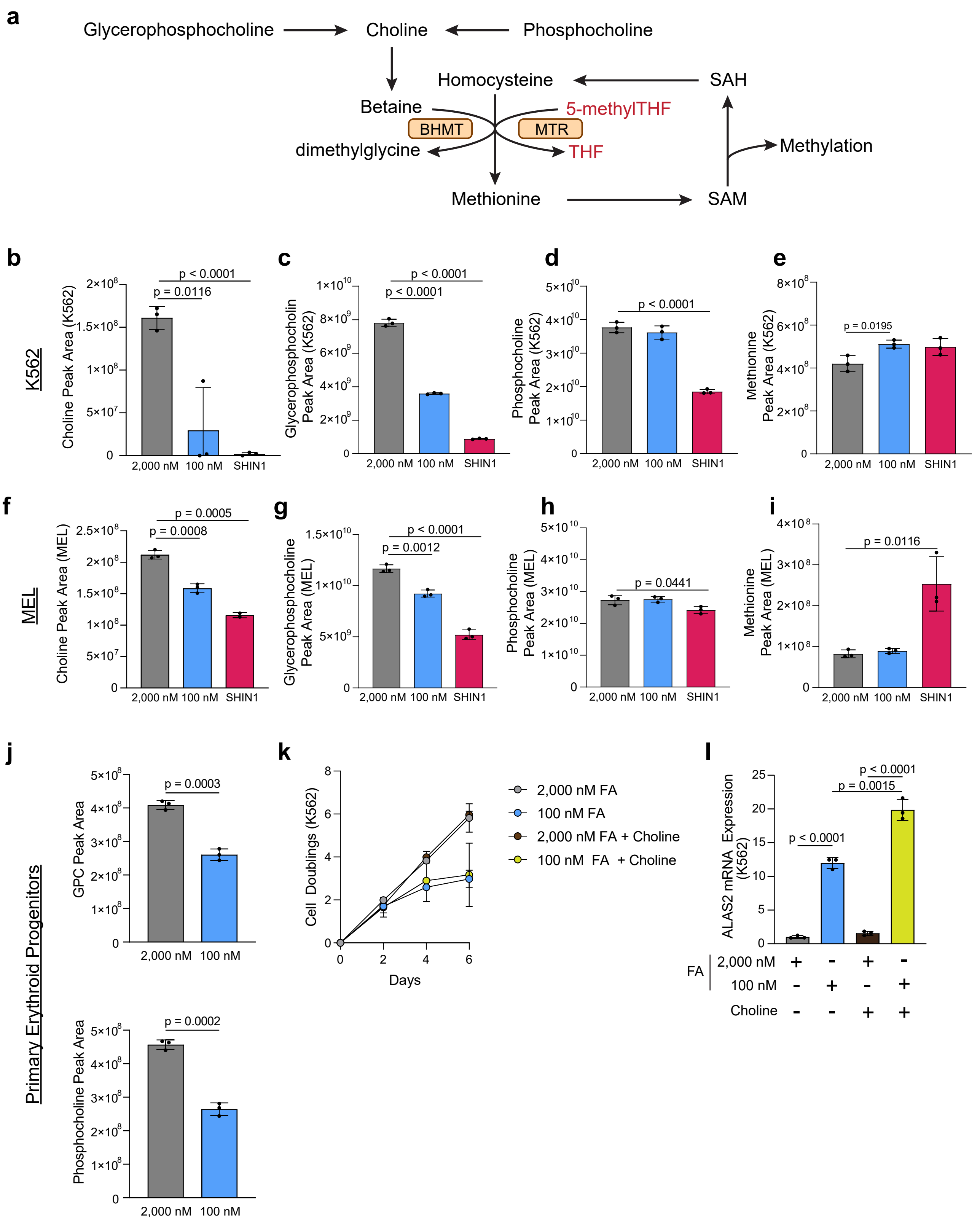

**Extended Data Figure 7.**

**a,** Schematic depicting the shared reaction between choline and folate metabolism. **b-e,** Indicated metabolite levels in K562 cultured for 2 days in 2,000 nM FA, 100 nM FA, and 2,000 nM FA + SHIN1 measured by LC-MS. **f-i,** Indicated metabolite levels measured by LC-MS in MEL in same conditions as b-e. **j,** glycerophosphocholine (GPC) and phosphocholine levels in murine primary erythroid progenitor cells cultured for 2 days in 2,000 and 100 nM FA. **k,** Cell proliferation of K562 in 2,000 and 100 nM FA with or without supplementation with choline (500 µM). **i,** RT-qPCR analysis of ALAS2 mRNA in choline-supplemented K562 cells following 6 days in 2,000 and 100 nM FA. Data shown are mean (± s.d.) of three biological replicates. All P values were calculated using an unpaired Student's t-test.

### Extended Data Figure 8

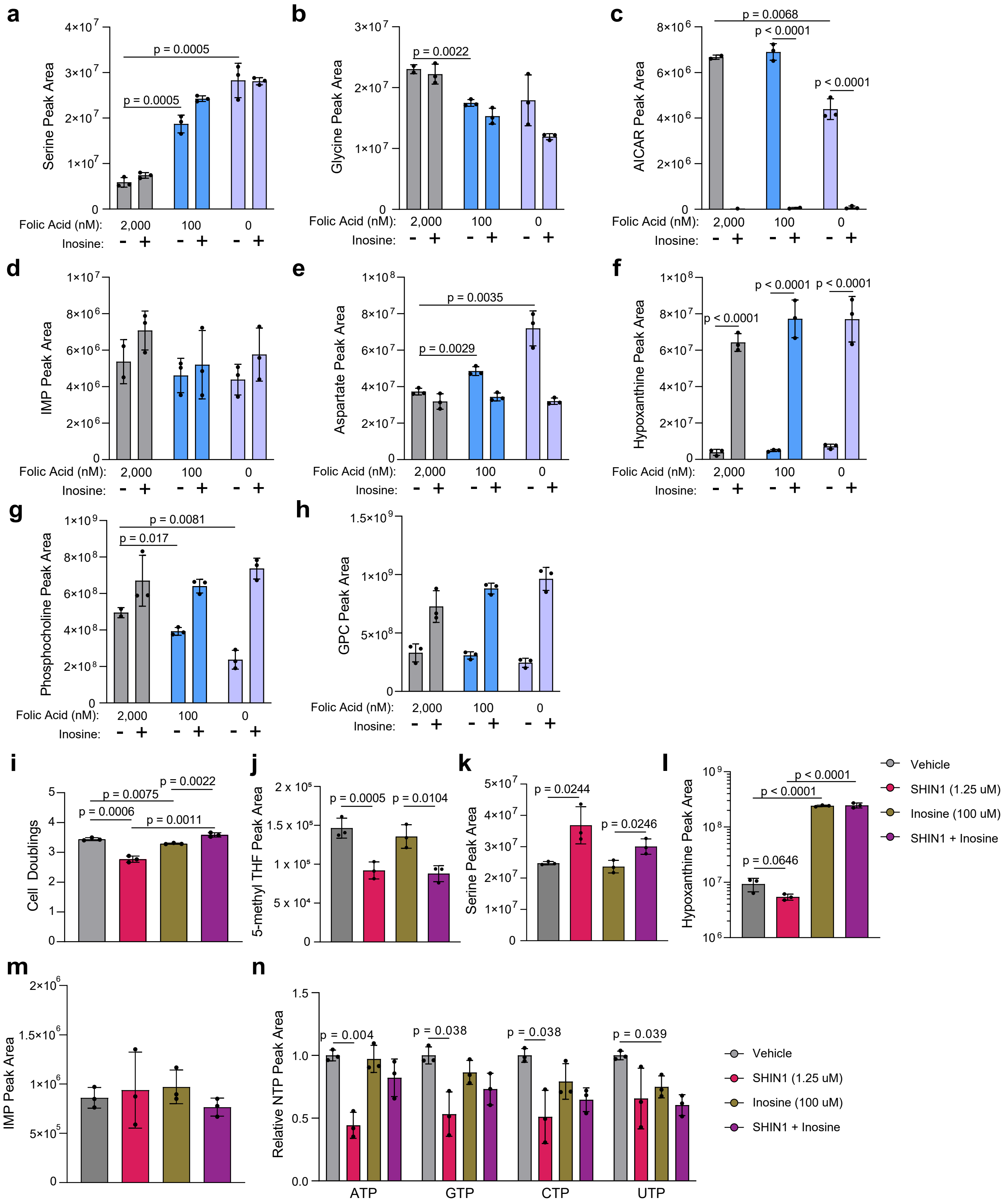

**Extended Data Figure 8.**

**a-h,** Indicated metabolite abundance as measured by LC-MS in murine primary erythroid progenitor cells cultured in 2,000, 100, or 0 nM FA for 2 days, with or without supplementation with inosine. **i,** Proliferation of vehicle-, SHIN1-, inosine-, or SHIN1 + inosine-treated murine primary erythroid progenitor cells (2 days), cultured in SFEM II expansion media. **j-n,** Indicated metabolite abundance measured by LC-MS in vehicle-, SHIN1-, inosine-, or SHIN1 + inosine-treated murine primary erythroid progenitor cells (2 days), cultured in SFEM II expansion media. GPC – glycerophosphocholine. Data shown are mean (± s.d.) of three biological replicates. All P values were calculated using an unpaired Student's t-test.

### Extended Data Figure 9

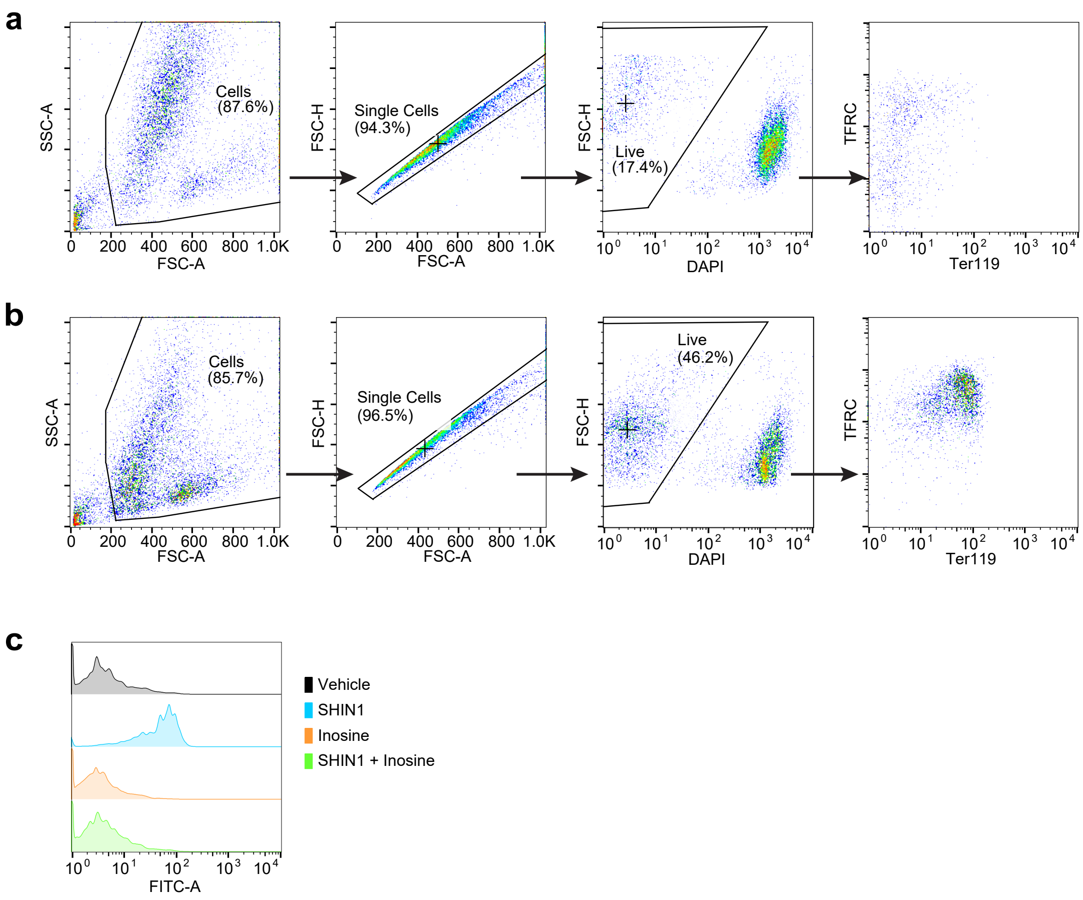

**Extended Data Figure 9.**

**a, b,** Representative flow cytometry populations showing erythroid progenitor cells in Vehicle (**a**) or SHIN1 (**b**) for 4 days. **c,** Representative histograms of Ter119 expression from erythroid progenitor cells cultured for 4 days in Vehicle, SHIN1, Inosine, or SHIN1 + Inosine.
